# Comparative Analysis of T Cell Spatial Proteomics and the Influence of HIV Expression

**DOI:** 10.1101/2021.03.17.435902

**Authors:** Aaron L. Oom, Charlotte A. Stoneham, Mary K. Lewinski, Alicia Richards, Jacob M. Wozniak, Km Shams-Ud-Doha, David J. Gonzalez, Nevan J. Krogan, John Guatelli

## Abstract

As systems biology approaches to virology have become more tractable, highly studied viruses such as HIV can now be analyzed in new, unbiased ways, including spatial proteomics. We employed here a differential centrifugation protocol to fractionate Jurkat T cells for proteomic analysis by mass spectrometry; these cells contain inducible HIV-1 genomes, enabling us to look for changes in the spatial proteome induced by viral gene expression. Using these proteomics data, we evaluated the merits of several reported machine learning pipelines for classification of the spatial proteome and identification of protein translocations. From these analyses we found that classifier performance in this system was organelle-dependent, with Bayesian t-augmented Gaussian mixture modeling outperforming support vector machine (SVM) learning for mitochondrial and ER proteins, but underperforming on cytosolic, nuclear, and plasma membrane proteins by QSep analysis. We also observed a generally higher performance for protein translocation identification using a Bayesian model, BANDLE, on SVM-classified data. Comparative BANDLE analysis of cells induced to express the wild-type viral genome vs. cells induced to express a genome unable to express the accessory protein Nef identified known Nef-dependent interactors such as TCR signaling components and coatomer complex. Lastly, we found that SVM classification showed higher consistency and was less sensitive to HIV-dependent noise. These findings illustrate important considerations for studies of the spatial proteome following viral infection or viral gene expression and provide a reference for future studies of HIV-gene-dropout viruses.

## Introduction

Spatial proteomics is a methodologically diverse and rapidly growing field within mass spectrometry (MS) that aims to understand the subcellular localization of the human proteome^1–7^. While initial efforts focused on establishing techniques and reference maps for various cell lines, recent work by the Cristea group expanded the field to understand the whole-cell effects of viral infection using human cytomegalovirus (HCMV) as a prototype^7^. This work led to novel findings on the importance of peroxisomes in herpesvirus infectivity^8^, exemplifying the power of these methods for uncovering new viral biology. However, as this was a first in its class study, how different methodologies might impact the results of viral studies using spatial proteomics is unclear. Using the well-characterized HIV-1 as a model virus system, we aimed to compare the output of several published spatial proteomic analysis pipelines^9–12^ as a survey of established methods.

To model HIV expression, we used a Jurkat T cell line that harbors a doxycycline-regulated HIV-1 genome. These cells were previously developed by our group to generate nearly homogenous HIV-positive cell populations for MS analysis^13^. As an additional biological comparator, we examined both wild-type (WT) virus and a virus lacking the accessory gene *nef* (ΔNef). Nef is a small (27 kDa), myristoylated membrane-associated accessory protein expressed early during the viral replication cycle^14, 15^. Nef increases viral growth-rate and infectivity^16^, and it dysregulates the trafficking of cellular membrane proteins such as CD4, class I MHC, and proteins involved in T cell activation such as CD28^17^ and p56-Lck^18^. Some of these activities enable the virus to evade immune detection^19, 20^. Here we use inducible Jurkat T cell lines containing either WT or ΔNef HIV-1NL4-3 provirus and compare the spatial proteome of uninduced cells to cells post-induction with doxycycline. To fractionate the cells, we used a modified version of the Dynamic Organellar Mapping protocol^5, 6^ with additional centrifugation steps^4^ to enhance organellar resolution, then analyzed the fractions by MS using TMT multiplexing.

Following the generation and processing of MS data, two broad steps are required for spatial proteomics: classification and hit determination. For classifying detected proteins into cellular organelles we compared two methods from pRoloc, an R software package developed by the Lilley lab^12^. The first was support vector machine (SVM) classification which outputs a label for each protein and an algorithm specific confidence score that can be used to threshold assignments^1^. The second was a Bayesian approach called t-augmented Gaussian mixture modeling with *maximum a posteriori* estimates (TAGM-MAP) which outputs a label for each protein and an actual probability of assignment^11^. To gauge the quality of these classifications, we compared the two methods using the QSep metric developed by the Lilley group^21^, which quantifies the separation, or resolution, of the organelles in question. We additionally cross-referenced our organellar assignments to existing organellar proteome databases^22–25^.

After classification, data were analyzed for translocating proteins following HIV expression. We compared three different methods for determining protein translocations: label-based movement, translocation analysis of spatial proteomics (TRANSPIRE)^9^, and Bayesian analysis of differential localization experiments (BANDLE)^10^. Label-based movement relies strictly on identifying proteins that are consistently classified in one organelle prior to a cellular perturbation, then consistently classified in another organelle following the perturbation; this method was employed by the Cristea group in their HCMV study^7^. TRANSPIRE is a refined methodology from the Cristea lab that relies on generating synthetic translocations from proteins of known localization and uses Bayesian analysis to determine the likelihood of proteins of unknown localization behaving in a manner consistent with anticipated translocations following a cellular perturbation^9^. Lastly, BANDLE is another method developed by the Lilley group that takes replicated data, both with and without a perturbation, and uses Bayesian analysis to yield a ranked list of possible translocations with their associated likelihood of occurrence^10^. We compared the hits from these various methods by cross-referencing hits with a previous study of the HIV interactome^26^ as well as the more broad NIH HIV-1 Human Interaction Database^27^.

From these comparisons we found that the performance of different classifiers is organelle-dependent and shows varied effects from HIV expression. As determined by agreement with previously published organellar proteomes, classification with TAGM-MAP showed increased accuracy in mitochondrial and ER-classified proteins, while SVM outperformed TAGM-MAP with nuclear, cytosolic, and plasma membrane-classified proteins. We also observed generally higher performance for protein translocation using BANDLE on SVM-classified data when compared to the HIV interactomes. BANDLE analysis of WT and ΔNef data identified known Nef interactors involved in T cell activation and the coatomer complex. Finally, we found that SVM classification showed higher consistency and was less sensitive to HIV-dependent noise. These findings illustrate the complexities in choosing a computational method for spatial proteomics study and serve as a foundation for additional studies.

## Experimental Procedures

### Experimental design and statistical rationale

All fractionation experiments with mass spectrometric analysis were performed in technical triplicate for each condition (uninduced and induced), with two biological replicates for wild-type and ΔNef NL4-3 Jurkat cells (Fig. 1A). This yielded a total of 6 uninduced and 6 induced technical replicates for each virus type. Biological replicates were prepared on separate days and analyzed by mass spectrometry on separate days. Western blotting and flow cytometry were performed on each technical replicate. Analyses for QSep (Fig. 2B and C) used Welch’s t-test to determine statistical significance.

**Figure 1:**
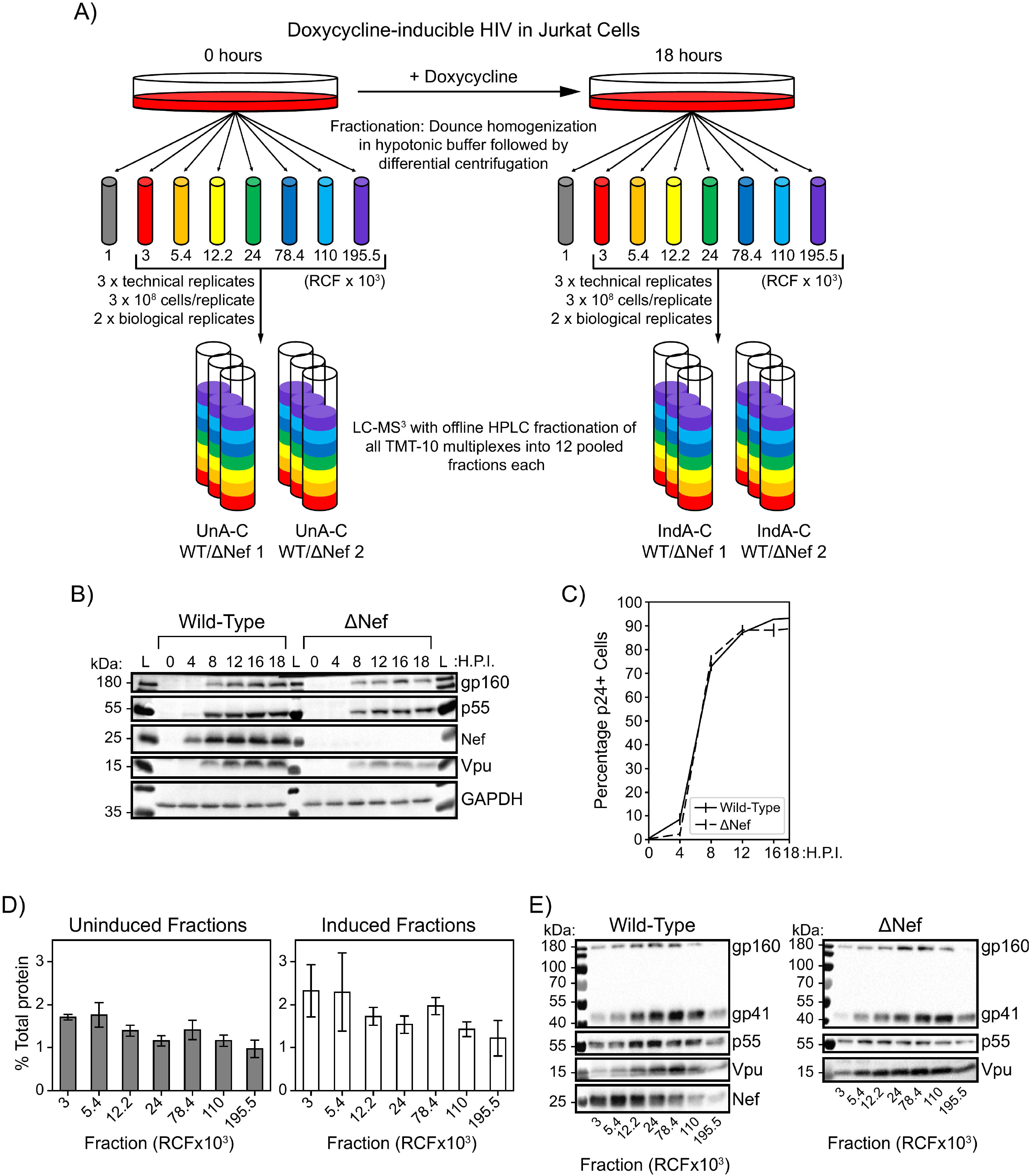
Inducible HIV-1 Jurkat cell lines yield a near pure population of HIV-expressing cells suitable for fractionation by differential centrifugation. A) Equal numbers of doxycycline-inducible wild-type and ΔNef HIV Jurkat cells were induced or left uninduced for 18 hours then fractionated by Dounce homogenization in a hypotonic lysis buffer. Cell homogenates were put through a differential centrifugation protocol, discarding the nuclear pellet (1,000x*g*) and lysing remaining pellets in 2.5% SDS buffer. Fractions were labeled for TMT-10 multiplexing and further offline HPLC fractionation. All multiplexes were run for 3 hours on LC-MS^3^. B) Western blot showing induction of HIV p55, gp160, gp41, Nef, and Vpu with a GAPDH loading control. Cells were induced for 0, 4, 8, 12, 16, and 18 hours, lysed, then a portion of these cell lysates was run on 10% SDS-PAGE gels. C) Flow cytometry analysis of remaining sample from 1B. HIV-1 expression peaked at ∼95% of cells p24+ by 18 hours. D) Average percentage of total cellular protein detected in each fraction by BCA protein assay. Bars represent the mean value for a given fraction based on the average from each biological replicate. Error bars are one standard deviation. All BCA assays were performed in technical triplicate on 10-fold dilutions for each biological replicate. E) Western blots for cell fractions of inducible wild-type HIV Jurkat cells (left) and ΔNef HIV Jurkat cells (right), 18 hours post-induction. Blots shown are representative of both biological replicates.

**Figure 2:**
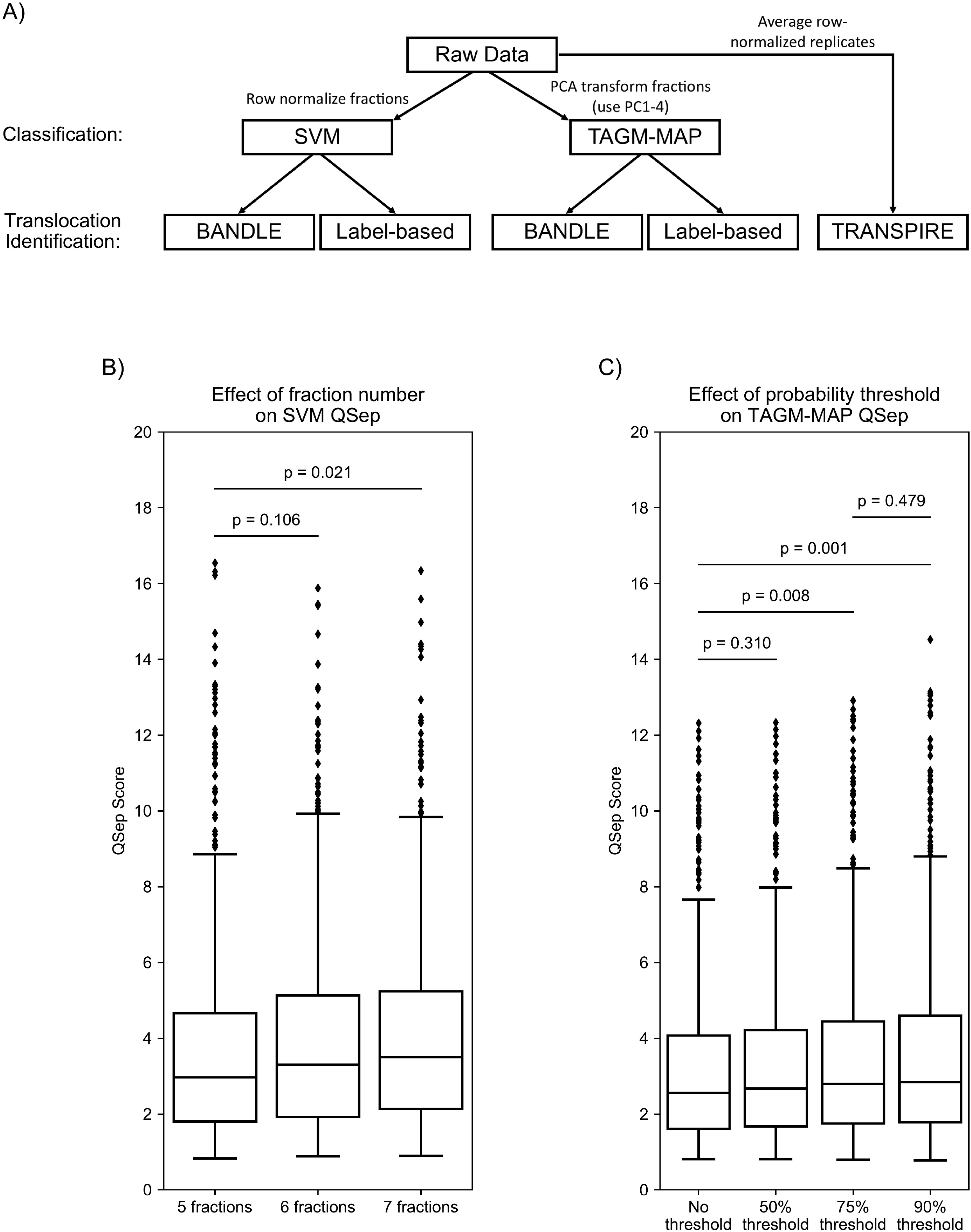
Analysis of fractionation data reveals increased organellar resolution from added fractions and thresholding TAGM-MAP data. A) Diagram of the computational methods used here. For SVM classification, the raw data of individual technical replicates were row normalized. For TAGM-MAP classification, the raw data of individual technical replicates were PCA transformed, with the first four principal components (PC1-4) carried forward for analysis. Both SVM and TAGM-MAP classified data were fed into BANDLE or label-based movement analysis. Lastly, for analysis with TRANSPIRE, individual technical replicates were row normalized then averaged together. B) Boxplot of QSep scores for SVM analysis of WT uninduced samples using the original 5 fractions described by Itzhak et al.^5^, adding a 110,000x*g* fraction (6 fractions), or adding both a 110,000x*g* and a 195,500x*g* fraction (7 fractions). C) Boxplot of QSep scores for TAGM-MAP analysis of WT uninduced samples comparing using no threshold for remaining classified, a 50% chance of classification, a 75% chance of classification, or a 90% chance of classification. Statistical significance is calculated using a two-sided, independent Student’s t-test with Welch’s correction for unequal variance. Boxplots show median, not mean, line.

### Cell culture

The doxycycline-inducible NL4-3 HIV-1 and NL4-3 ΔNef Jurkat cell lines were previously described^13, 28^. The replication-incompetent genome used was based on pNL4-3 but lacked most of the 5’ U3 region, encoded a self-inactivation deletion in the 3’ LTR, and contained the V3 region from the R5- tropic 51-9 virus^29^ to prevent the cell-cell fusion of the Jurkat T cells used herein, which do not express CCR5. Inducible cells were cultured in RPMI 1640 media supplemented with penicillin/streptomycin (pen/strep) and 10% Tet-free fetal bovine serum (FBS), as well as puromycin (1 µg/mL) and G418 (200 µg/mL) to maintain persistence of the tetracycline trans-activator and the inducible genome. Cells were passaged every two days to keep concentrations between 3.5×10^5^ and 1×10^6^ cells/mL. Cells were maintained at 37°C, 5% CO2, and 95% humidity.

### Doxycycline induction and fractionation

On the day before fractionation, 2.016×10^9^ cells were plated at 6×10^5^ cells/mL in T75 flasks at a total volume of 40 mL/flask. Half of these cells were induced to express HIV-1/HIV-1ΔNef with doxycycline (1 µg/mL) for 18 hours, while the other half remained uninduced. Following induction, cells of each condition, i.e. uninduced and induced, were split into three technical replicates, and then centrifuged at 500x*g* for 5 min at 4°C. Each technical replicate was pooled into a single 50 mL tube using ice cold 1X PBS, then counted by hemocytometer. From each technical replicate, 3×10^8^ cells were fractionated. Two aliquots of cells were taken from each technical replicate for whole cell western blots and testing induction by flow cytometry.

The fractionation protocol used here is derived from the Dynamic Organellar Maps method^5^ with additional centrifugation steps^4^ and TMT-based MS analysis rather than SILAC^6^. Cells for fractionation were centrifuged at 500x*g* for 5 min at 4°C then resuspended in ice-cold PBS and incubated for 5 min on ice. Cells were again centrifuged at 500x*g* for 5 min at 4°C, then resuspended in ice-cold hypotonic lysis buffer (25 mM Tris-HCl (pH 7.5), 50 mM sucrose, 0.5 mM MgCl2, and 0.2 mM EGTA in water) and incubated for 5 min on ice. Using a 7 mL Dounce homogenizer, cells were homogenized with 20 full strokes of the tight pestle. Cell homogenates were then immediately transferred to a 13 mL (14×89 mm) ultracentrifuge tube with sufficient ice-cold hypertonic sucrose buffer (1.25 M sucrose, 25 mM Tris-HCl (pH 7.5), 0.5 mM MgCl2, and 0.2 mM EGTA in water) to restore 250 mM sucrose concentration. All replicates were then centrifuged at 1,000x*g* for 10 min at 4°C in a Beckman Coulter ultracentrifuge (SW- 41 Ti rotor), balancing each tube with balance buffer (250 mM sucrose, 25 mM Tris-HCl (pH 7.5), 0.5 mM MgCl2, and 0.2 mM EGTA in water). Supernatants were transferred to a fresh ultracentrifuge tube, balanced with balance buffer, then fractionated using the following differential centrifugation protocol: 3,000x*g* for 10 min, 5,400x*g* for 15 min, 12,200x*g* for 20 min, 24,000x*g* for 20 min, 78,400x*g* for 30 min, 110,000x*g* for 35 min, and 195,500x*g* for 40 min All centrifugation steps were performed at 4°C with pellets from each spin being resuspended in SDS buffer (2.5% SDS and 50 mM Tris-HCl (pH 8.0) in water). Fractions were then heated for 10 minutes at 72°C. Protein content of each fraction was quantified in triplicate using a bicinchoninic acid (BCA) protein assay (Thermo-Fisher).

### Confirmatory western blots and p24 flow cytometry

Prior to mass spectrometric analysis of fractions, induction and fractionation were evaluated by flow cytometry and western blotting (Fig. 1B and C). For p24 flow cytometry, an aliquot of 2×10^6^ cells from each technical replicate were pelleted at 500x*g* for 5 min at 4°C then resuspended in ice-cold FACS buffer (2% FBS and 0.1% sodium azide in 1X PBS). The cells were again pelleted at 500x*g* for 5 min at 4°C then resuspended in Cytofix/Cytoperm reagent (BD Biosciences) and incubated on ice for 30 min Following fixation/permeabilization, cell suspensions were diluted with wash buffer and pelleted at 500x*g* for 5 min at 4°C. Cells were resuspended in p24 primary antibody solution (1:100 dilution of p24-FITC antibody clone KC57 (Beckman Coulter) diluted in perm/wash buffer) and incubated on ice for 30 min in darkness. Ice-cold FACS buffer was added to each sample and cells were pelleted at 500x*g* for 5 min at 4°C. The intracellular p24 was analyzed using an Accuri C6 flow cytometer (BD Biosciences). Uninduced cells had an average p24+ population of 0.27% (S.D. = 0.20) and live cell population of 85.78% (S.D. = 3.37). Induced cells had an average p24+ population of 94.85% (S.D. = 1.23) and live cell population of 79.25% (S.D. = 4.35).

An aliquot of 1×10^7^ cells from each technical replicate was lysed in SDS buffer and probe sonicated on ice until no longer viscous. 3,000x*g* fractions were also probe sonicated. The samples were mixed with 4X loading buffer (200 mM Tris-HCl (pH 6.8), 8% SDS, 40% glycerol, 200 mM tris(2- carboxyethyl)phosphine-HCl (TCEP), and 0.04% bromophenol blue in water) and proteins were then separated on 10% SDS-PAGE gels at a constant 70V. Proteins were transferred to polyvinylidene difluoride (PVDF) membranes for 1 hour using the Trans-Blot turbo (BioRad) system using standard conditions. The membranes were blocked in 5% milk in 1X PBS-T for 30 min at room temperature prior to incubation with primary antibodies diluted in 1% milk and 0.05% sodium azide in 1X PBS-T: sheep anti-Nef (gift from Celsa Spina, diluted 1:3,000), mouse anti-p24 (Millipore, diluted 1:500), Chessie8 (mouse anti-gp41, NIH AIDS Research and Reference Reagent program^30^, diluted 1:10,000), rabbit anti-Vpu (NIH AIDS Research and Reference Reagent program ARP-969, contributed by Dr. Klaus Strebel, diluted 1:1,000), and mouse anti-GAPDH (GeneTex, diluted 1:5,000). The blots were washed and probed with either horseradish peroxidase-conjugated goat anti-mouse, HRP-goat anti-rabbit, or HRP-rabbit anti-sheep secondary (BioRad) diluted 1:3,000, incubating for 1 hour at room temperature on a shaker. Apparent molecular mass was estimated with PageRuler protein standard (Thermo Scientific). Blots were imaged using Western Clarity detection reagent (BioRad) before detection on a BioRad Chemi Doc imaging system with BioRad Image Lab v5.1 software.

### Sample digestion for mass spectrometry

Disulfide bonds were reduced with 5 mM TCEP at 30°C for 60 min and cysteines were subsequently alkylated (carbamidomethylated) with 15 mM iodoacetamide (IAA) in the dark at room temperature for 30 min Proteins were then precipitated with 9 volumes of methanol, pelleted and resuspended in 1M urea, 50 mM ammonium bicarbonate. Following precipitation, protein concentration was determined using a BCA protein assay. A total of 0.2 mg of protein was subjected to overnight digestion with 8.0 µg of mass spec grade Trypsin/Lys-C mix (Promega). Following digestion, samples were acidified with formic acid (FA) and subsequently 150 ug peptides were desalted using AssayMap C18 cartridges mounted on an Agilent AssayMap BRAVO liquid handling system, C18 cartridges were first conditioned with 100% acetonitrile (ACN), followed by 0.1% FA. The samples were then loaded onto the conditioned C18 cartridge, washed with 0.1% FA, and eluted with 60% MeCN, 0.1% FA. Finally, the organic solvent was removed in a SpeedVac concentrator prior to LC-MS/MS analysis.

### TMT Labeling

Peptide concentration was determined using a Nanodrop, and a total of 15 µg of peptide was then used for TMT labeling, each replicate serving as a multiplex. Briefly, dried peptide sample was resuspended in 200 mM HEPES (pH 8) and incubated for 1 h at room temperature with one of the TMT10-plex reagents (ThermoFisher) solubilized in 100% anhydrous ACN. Reactions were quenched using a 5% hydroxylamine solution at 1-2 μl per 20 μl TMT reagent. The multiplexed samples were then pooled and dried in a SpeedVac. The labeled peptides were resuspended in 0.1% FA. After sonication for 1min, the sample was desalted manually using SepPak; the column was first conditioned with 100% ACN, followed by 0.1% FA. Sample was loaded, then washed with 0.1% FA and eluted in a new vial with 60% ACN, 0.1% FA. Finally, the organic solvent was removed using a SpeedVac concentrator prior to fractionation.

### High pH Reverse-Phase Fractionation

Dried samples were reconstituted in 20mM ammonium formate (pH ∼10) and fractionated using a Waters ACQUITY CSH C18 1.7 μm 2.1 × 150 mm column mounted on a MClass Ultra Performance Liquid Chromatography (UPLC) system (Waters corp., Milford, MA) at a flow rate of 40 μl/min with buffer A (20 mM ammonium formate pH 10) and buffer B (100% ACN). Absorbance values at 215 nm and 280 nm were measured on a Waters UV/Vis spectrophotometer, using a flowcell with a 10 mm path length. Peptides were separated by a linear gradient from 5% B to 25% B in 62.5 min followed by a linear increase to 60% B in 4.5 min and 70% in 3 min and maintained for 7 min before increasing to 5% in 1 min Twenty-four fractions were collected and pooled in a non-contiguous manner into twelve total fractions. Pooled fractions were dried to completeness in a SpeedVac concentrator.

### LC-MS^3^ Analysis

Dried samples were reconstituted with 0.1% FA and analyzed by LC-MS/MS on an Orbitrap Fusion Lumos mass spectrometer (Thermo) equipped with an Easy nLC 1200 ultra-high pressure liquid chromatography system interfaced via a Nanospray Flex nanoelectrospray source (Thermo). Samples were injected on a C18 reverse phase column (25 cm x 75 um packed with Waters BEH 1.7 um particles) and separated over a 120-min linear gradient of 2-28% solvent B at a flow rate of 300nL/min The mass spectrometer was operated in positive data-dependent acquisition mode.

Parameter settings were set as follows: FT MS1 resolution (120 000) with AGC target of 1e6, ITMS2 isolation window (0.4 m/z), IT MS2 max. inject time (120 ms), IT MS2 AGC (2E4), IT MS2 CID energy (35%), SPS ion count (up to 10), FT MS3 isolation window (0.4 m/z), FT MS3 max. inject time (150 ms), FT MS3 resolution (50 000) with AGC target of 1e5. A TOP10 method was used where each FT MS1 scan was used to select up to 10 precursors for interrogation by CID MS2 with readout in the ion trap. Each MS2 was used to select precursors (SPS ions) for the MS3 scan which measured reporter ion abundance.

### Mass spectrometry spectra identification

Raw files were analyzed using Proteome Discoverer v2.3 (Thermo Fisher Scientific). MS/MS spectra were searched against a concatenated database containing Uniprot human and HIV-1 proteins (downloaded 02/03/20) and reverse decoy sequences using the Sequest algorithm^31^; the database contained 20,367 total entries. Mass tolerance was specified at 50 ppm for precursor ions and 0.6 Da for MS/MS fragments. Static modifications of TMT 10-plex tags on lysine and peptide n-termini (+229.162932 Da) and carbamidomethylation of cysteines (+57.02146 Da), and variable oxidation of methionine (+15.99492 Da) were specified in the search parameters. Data were filtered to a 1% false discovery rate at the peptide and protein level through Percolator^32^ using the target-decoy strategy^33^. TMT reporter ion intensities were extracted from MS3 spectra within Proteome Discoverer to perform quantitative analysis.

### Computational analysis

Matching biological replicates were combined (i.e. WT biological replicate 1 and 2), then analyzed using the various pipelines described. The *Homo sapiens* (“hsap”) marker set from pRoloc was used in all cases. For classification and hit generation, only the proteins commonly detected across matched biological replicates were analyzed to allow for consistency in comparing methods on the same data set.

The pRoloc implementation of SVM^12^ was performed on row-normalized data sets, while the pRoloc implementation of TAGM-MAP^11^ required PCA transformation and no row-normalization with the first four principal components carried forward. The PCA transformation was used because of floating point arithmetic errors that arose because of highly correlated features. Default parameters for algorithms were used excepting the following:

> SVM hyperparameter classification: 10 times 10-fold cross-validation
>
> SVM classification threshold: median algorithm score for each organelle
>
> TAGM-MAP model training: 200 iterations
>
> BANDLE: 6 chains

TRANSPIRE was run on averaged row-normalized datasets, i.e., technical replicates were row-normalized then values for each feature were averaged for each protein across matched technical replicates. Organelles were combined into 5 groups: 1) Golgi apparatus/plasma membrane/endoplasmic reticulum/peroxisomes/lysosomes, 2) cytosol/actin cytoskeleton/proteasome, 3) nucleus, 4) mitochondria, and 5) 40S/60S ribosome. The number of inducing points and the kernel function were chosen from amongst the suggested values in the TRANSPIRE documentation. For these datasets, 75 inducing points and the squared exponential kernel performed best and were used in the analysis.

The average distribution of proteins across organelles was calculated by determining the average organellar distribution for a single technical replicate, then averaging the values of matched technical replicates. Marker profiles were generated by averaging the behavior of markers for a given organelle within a technical replicate, then averaging those values across technical replicates for each organelle. Organellar QSep scores were calculated by averaging the individual QSep scores between two organelles across all matched technical replicates, then plotting the distribution of those averages.

Comparisons to the Human Protein Atlas (HPA) were completed by combining several HPA subcellular localization annotations to align with the organelles used by pRoloc:

1. Nuclear membrane, nucleoli fibrillar center, nucleoli rim, nucleoli, kinetochore, mitotic chromosome, nuclear bodies, nuclear speckles, and nucleoplasm: Nucleus
2. Actin filaments and focal adhesion sites: Actin Cytoskeleton
3. Plasma membrane and cell junctions: Plasma Membrane

Remaining designations within the HPA beyond the above and those in common with pRoloc’s “hsap” markers were not considered. The 40S Ribosome, 60S Ribosome, and Proteasome classes from the SVM and TAGM-MAP classified data were collapsed into the Cytosol label.

Thresholds for Figures 6 and S8 were determined by dividing the size of the Jäger HIV interactome^26^, 453 proteins, or the NIH HIV interactome^27^, 4,628 proteins, by the predicted human proteome size of 19,773 proteins^34^. G.O. analysis for Figure 6B was conducted using the STRING database^35^.

## Results

### Doxycycline-inducible HIV-1NL4-3 Jurkat T cells are a scalable and uniform system for subcellular fractionation and proteomic studies

The WT HIV-1 inducible cells used here were previously generated and used for whole-cell quantitative- and phospho-proteomics^13^. To avoid the formation of syncytia, which could alter the subcellular fractionation and subsequent spatial proteomic data, the inducible HIV-1NL4-3 genomes were modified with a CCR5-tropic Env protein to avoid cell-cell fusion between the CCR5-negative Jurkat cells. Due to the high induction rates of HIV-1 expression and the scalability of this culture system, we reasoned that it would be amenable to subcellular fractionation by differential centrifugation with subsequent MS analysis (Fig. 1A). To determine the optimal time-point for analysis following induction of HIV-1 expression, cells were treated with doxycycline for 0, 4, 8, 12, 16, and 18 hours, and the expression of HIV-1 proteins was detected by western blotting and flow cytometry (Fig. 1B and C). WT cells began to express detectable Nef by 4 hours post-induction, and both WT and ΔNef cells expressed p55 Gag precursor (the precursor protein for virion structural proteins) by 8 hours and gp160 (the envelope glycoprotein precursor) by 12 hours. By 18 hours, viral proteins were robustly expressed; about 90-95% of both WT and ΔNef cells were positive by flow cytometry for p24 capsid (a proteolytic product of p55).

Subcellular fractionation was performed 18 hours post-induction; the cells were mechanically ruptured with a Dounce homogenizer in hypotonic solution, then subjected to a differential centrifugation protocol before preparation for quantitative, multiplexed MS analysis. Uninduced and induced cells were handled in technical triplicate for each biological replicate (n=2). We used a modified version of the Dynamic Organellar Mapping (D.O.M.) protocol^5, 6^ with additional fractions generated at 110,000x*g* and 195,500x*g* to increase the resolution of the classification analysis; a similar method of expanded differential centrifugation fractionation was previously described by the Lilley group^36^. As a quality control before MS, protein yields were quantified for each fraction (Fig. 1D). The post-nuclear fractions accounted for only ∼10-15% of total cellular protein, presumably because nuclear proteins and soluble cytoplasmic proteins that failed to pellet at 195,500x*g* were discarded, leaving primarily membranous organelles or organellar fragments and large, cytoplasmic complex proteins in the fractions analyzed. We also observed decreasing protein yields across the fractions, with an increase in the 78,400x*g* fraction, consistent with the original D.O.M. study using HeLa cells^5^. In further support of differential fractionation, varied abundances of viral proteins across the fractions in cells expressing either the WT or ΔNef genomes were observed by western blotting (Fig. 1E). Following confirmation of differential fractionation, we analyzed all fractions by LC-MS^3^ with TMT-10 multiplexing (Fig. 1A).

To determine the consistency of the MS analysis we used unsupervised hierarchical clustering by Spearman correlation coefficient for the individual fractions. We found that for both the WT and ΔNef data the fractions clustered by *g*-force rather than biological replicate (Fig. S1 and 2), suggesting consistent quantification values. Because the WT and ΔNef Jurkat cell lines represent individual clones for each, we also compared the uninduced fractions of the WT and ΔNef data to each other. This comparison showed that fractions still clustered by *g*-force rather than HIV genome (Fig. S3).

### SVM shows greater organellar resolution than TAGM-MAP even with stringent thresholds of classification for TAGM-MAP

To classify the fractionation data and identify translocating proteins, we employed a variety of previously published methods (Fig. 2A). As several resources detail known HIV interactors^26, 27^, we primarily focused on comparing classification and translocation identification methods using our WT data. In subsequent analyses, we examined the ΔNef data to determine the power of various methods in identifying Nef-specific effects.

For classification, proteins were classified using either the pRoloc implementation of SVM or TAGM-MAP. As the differential centrifugation protocol employed here is a modified version of the D.O.M. method which generates only 5 fractions^5^, we first examined whether our two additional fractions improved organellar resolution. The D.O.M. method classifies proteins with SVM, so we compared the resolution of organelles with the QSep analysis^21^ using the first 5 fractions for SVM classification, then the first 6 fractions, and finally all 7 fractions (Fig. 2B). We found that while the addition of the 110,000x*g* spin alone had no significant effect on organellar resolution as compared to the original method, the subsequent addition of the 195,500x*g* spin yielded a significant increase from a mean QSep score of 3.74 to 4.05 (median scores 2.97 and 3.50, respectively). In light of this, all subsequent analyses on the SVM data were performed on the full 7 fractions.

To determine if an alternate method for classification would perform better than SVM, we also tested the pRoloc implementation of TAGM-MAP. The outputs from TAGM-MAP give both a localization and a probability that the given protein is located in that organelle. These probabilities allowed us to test the effect of different probability thresholds on TAGM-MAP’s QSep scores. While using a 50% threshold, i.e. converting all proteins with a probability of localization lower than 50% to an “unknown” designation, showed no significant effect, 75% and 90% thresholds both showed significant gains over no thresholding (Fig. 2C). A 90% threshold showed no significant increase in QSep scores over the 75% threshold, so subsequent analyses employed the 75% threshold for TAGM-MAP classification. Of importance, we observed that the QSep scores from SVM classification were on average higher than those from TAGM-MAP even when comparing TAGM-MAP’s highest condition (90% probability threshold, average score of 3.55) to SVM’s lowest condition (5 fractions, average score of 3.74).

### SVM classifies proteins more consistently than TAGM-MAP

We next wanted to understand how the SVM and TAGM-MAP methods compared for consistency of classification across WT replicates (Fig. 3A and B). Both SVM (Fig. 3A) and TAGM-MAP (Fig. 3B) showed a low percentage (∼10-15%) of proteins that were classified identically in 6 out of 6 technical replicates for either WT uninduced or induced. However, allowing for a majority of replicates, i.e. 4 out of 6, gave ∼70-75% of proteins as classified consistently by SVM (Fig. 3A). This compared to ∼50-55% of proteins classified to a similar consistency by TAGM-MAP (Fig. 3B). HIV expression modestly decreased the consistency of both SVM and TAGM-MAP (∼5% difference), suggesting an increase in experimental noise from HIV expression.

**Figure 3:**
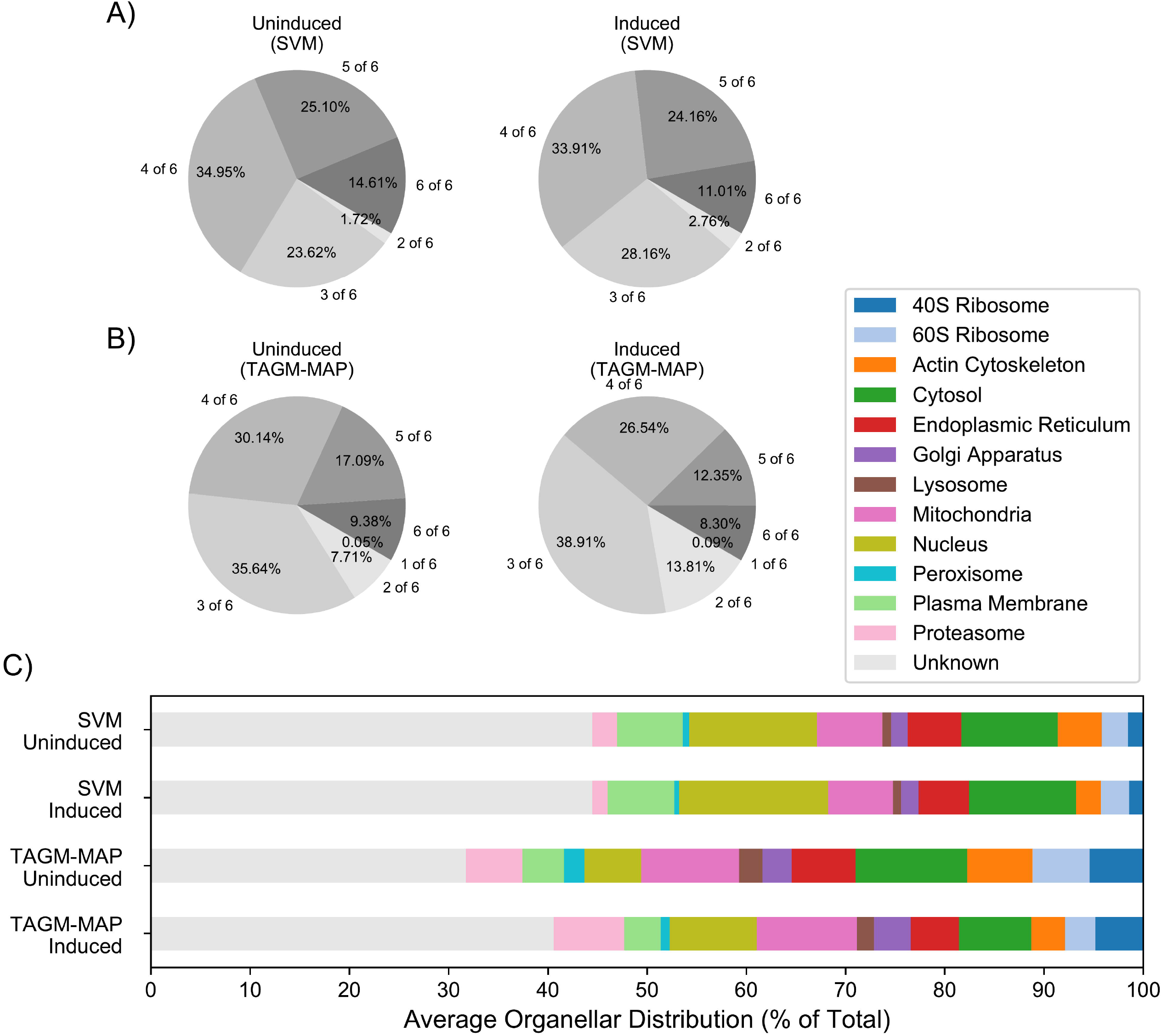
Classification with SVM shows greater consistency than TAGM-MAP classification. A) Proteins were classified by SVM and the most frequent organellar classification was identified along with its frequency, i.e. number of technical replicates classified as such. Left pie chart shows consistency of classification for WT uninduced replicates and right pie chart shows WT induced replicates. B) Same as A), but classification by TAGM-MAP. C) Average distribution of proteins across organelles for each indicated condition. All charts consider the same common proteins found across all WT replicates (4,765 proteins).

Looking at the average distribution of proteins across organelles, we found that SVM yielded a higher percentage of proteins that reverted to an unknown designation (Fig. 3C, 44% of proteins); this may partly explain the higher QSep scores generally seen for SVM compared to TAGM-MAP (Fig. 2). However, this percentage is stable between WT uninduced and induced replicates, while the lower percentage of unknown proteins (32% for uninduced and 41% for induced) for TAGM-MAP is more sensitive to HIV expression. Similar trends were seen within the ΔNef data (Fig. S4); marker behavior for WT (Fig. S5) and ΔNef (Fig. S6) is also similar, which likely explains the consistent trends. These data show a greater consistency for SVM classification and additionally suggest that SVM is less susceptible to noise introduced into data by HIV expression.

### Agreement between SVM and TAGM-MAP classification is organelle-dependent and is variably affected by HIV expression

To determine the concordance of SVM and TAGM-MAP for classification, we examined all proteins that were classified consistently in at least 4 of 6 WT replicates for both SVM and TAGM-MAP. We found more such proteins for the uninduced replicates (Fig. 4A), 1,863 proteins, as compared to the induced replicates (Fig. 4B) with 1,448 proteins. This difference may be attributable to the decrease in classification consistency caused by HIV expression for both SVM and TAGM-MAP, which would be accentuated by any increased susceptibility of TAGM-MAP to HIV-dependent noise. Of these consistently classified proteins, HIV expression minimally affected classifier agreement; 65% agreed between SVM and TAGM-MAP for WT uninduced and 69% agreed between SVM and TAGM-MAP for induced replicates (see diagonal of heatmaps). However, HIV expression increased the proportion of proteins that were consistently designated unknown by both SVM and TAGM-MAP: in uninduced cells, 40% of proteins agreed upon by the two methods were designated unknown (Fig. 4A), while 71% of agreed upon proteins were designated unknown from induced cells (Fig. 4B). This shift seems primarily driven by the increase in unknown designations for TAGM-MAP following HIV expression: in uninduced replicates, 52% of proteins designated unknown by SVM agreed with TAGM-MAP, but in induced replicates, 81% of these proteins agreed with TAGM-MAP. Matching trends were seen in ΔNef data (Fig. S7). Taken together, these data suggest that while HIV expression has little effect on the proportion of consistently classified proteins that are agreed upon by the two classifiers, the proportion of these proteins that are designated unknown is increased, and the overall number of consistently classified proteins is decreased.

**Figure 4:**
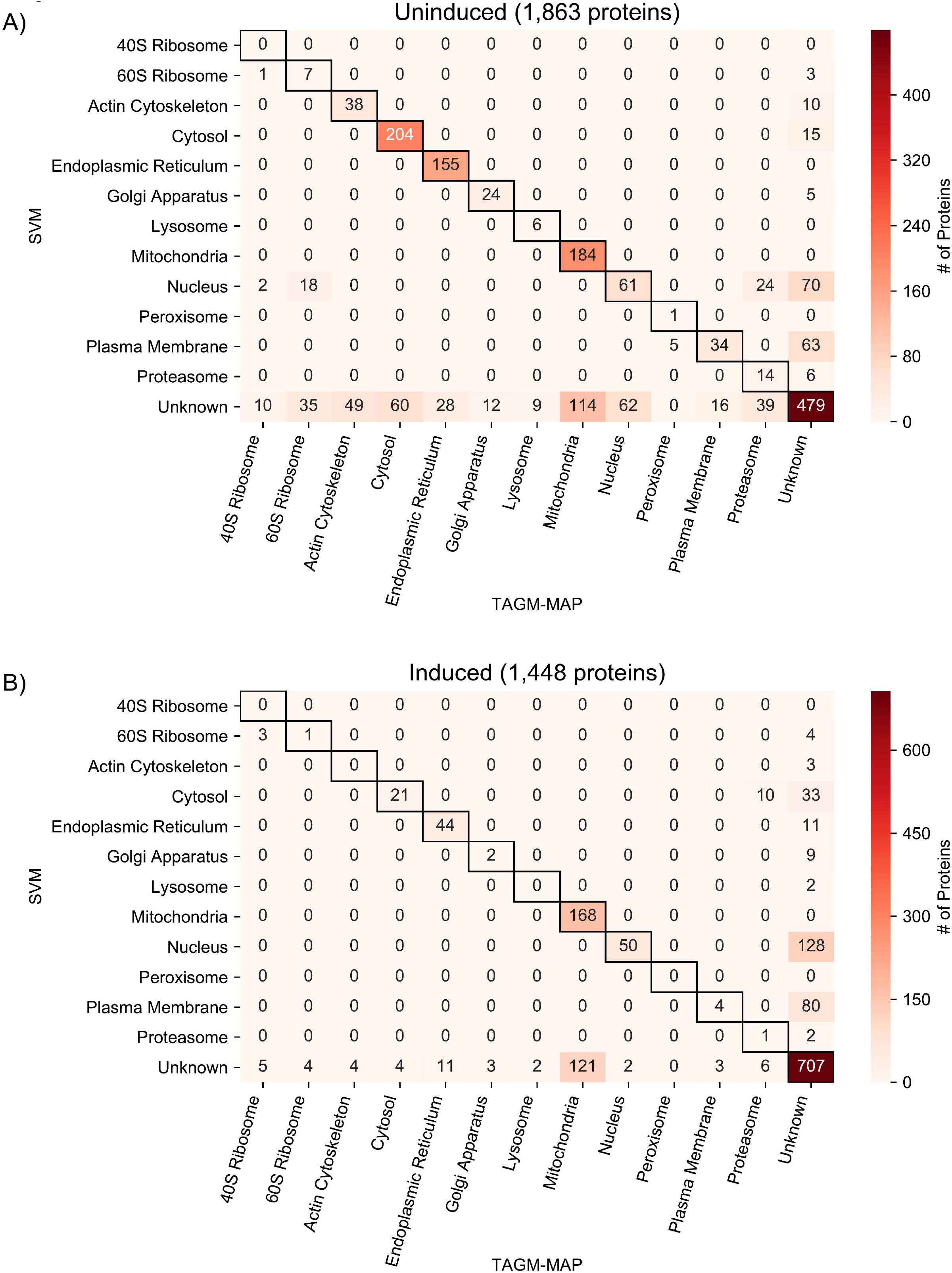
Concordance of SVM and TAGM-MAP classifications depends on organelle and expression of HIV. A) Heat map of common proteins that were consistently classified (proteins classified consistently in at least 4 of 6 replicates) by both SVM and TAGM-MAP for uninduced condition. Annotations indicate number of proteins in a given scenario. B) Same as A) for induced condition.

We found that proteins from the cytosol, ER, and mitochondria were the most frequent among consistently classified proteins. These three organelles also showed the best agreement between SVM and TAGM-MAP for uninduced replicates (Fig. 4A and S4A). However, HIV expression decreased the proportion of cytosolic proteins and ER proteins in agreement between SVM and TAGM-MAP: 73% of all proteins classified as cytosolic and 85% of all proteins classified as ER agreed for WT uninduced replicates, but only 31% of cytosolic proteins and 67% of ER proteins agreed for induced replicates. This decrease was smaller for mitochondrial proteins: 62% for uninduced and 58% for induced. Similar trends for cytosolic and mitochondrial proteins were seen in ΔNef data, but ER proteins showed little change (Fig. S7). These data show an organelle-dependent trend in classifier agreement that is variably affected by HIV expression.

### TAGM-MAP classification yields higher agreement than SVM classification with reported ER and mitochondria proteomes, but lower agreement in other organelles

To gauge the quality of our classifications, we compared those proteins that were consistently classified, i.e. 4 out of 6 replicates, for WT uninduced to several published spatial proteomes: MitoCarta2.0 database^22^, a study of the mitochondrial matrix proteome^23^, and a review of lysosome proteomic studies^24^ (Fig. 5A). Examining those proteins from each study that were detected in our datasets, we found that TAGM-MAP consistently out-performed SVM for mitochondria but performed less well for lysosomes. We also compared only those proteins that received an organellar classification, i.e. we excluded consensus unknown designations, to see if a focus on only proteins that remained classified would change the performance of SVM (orange bars) or TAGM-MAP (dark orange bars). SVM was more responsive to the exclusion of unknown proteins compared to TAGM-MAP, which is likely due to the lower proportion of unknown proteins in the TAGM-MAP uninduced condition.

**Figure 5:**
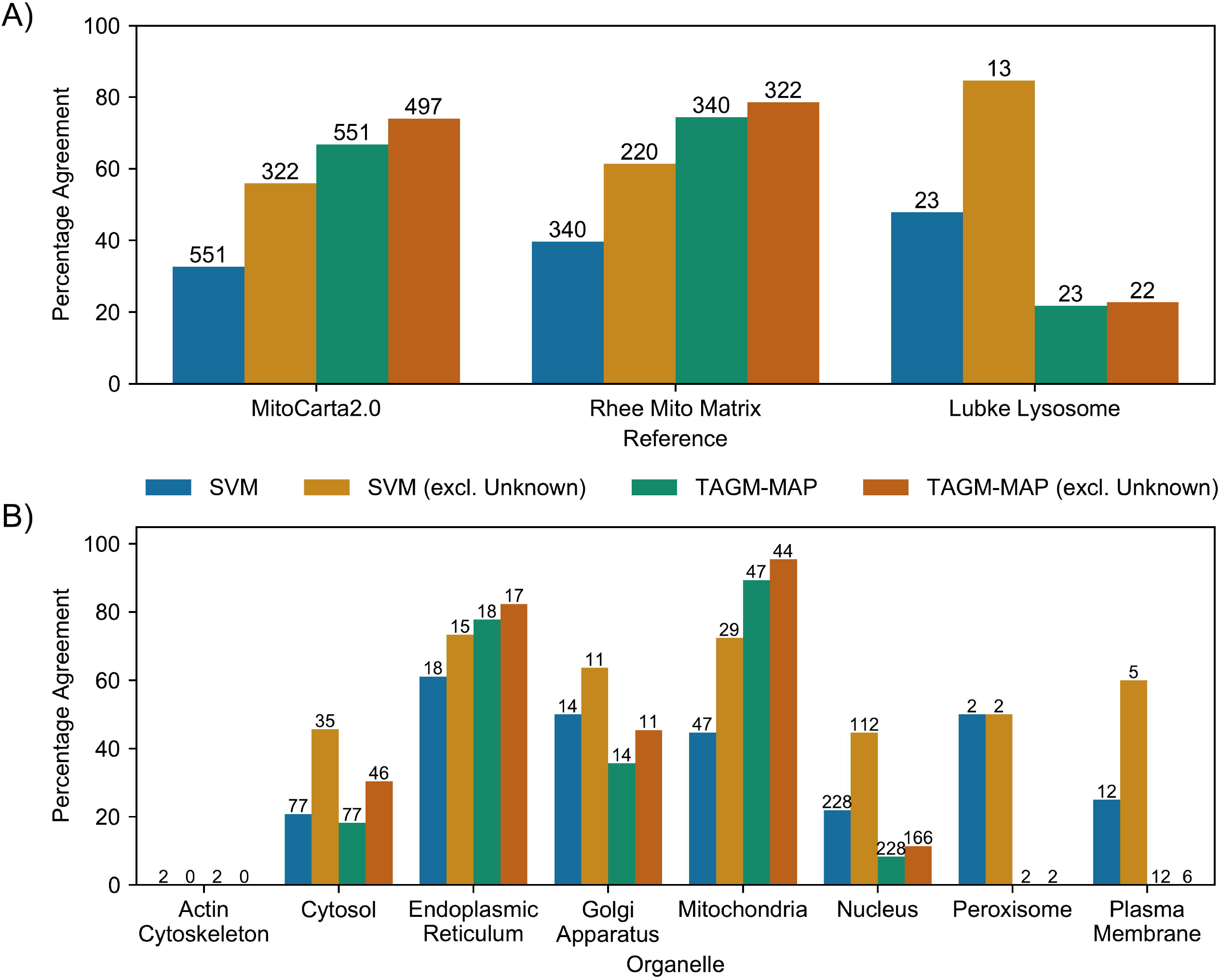
Validation of protein classification reveals higher performance for ER and mitochondria using TAGM-MAP, but better performance for Golgi apparatus, nucleus, and plasma membrane using SVM. A) Percentage of detected proteins from MitoCarta2.0 database^22^, Rhee et al. mitochondrial matrix study^23^, or Lubke lysosome proteome^24^ that were consistently classified (proteins classified consistently in at least 4 of 6 replicates) in line with the respective reference. Numbers above bars indicate the total number of proteins from that reference that were detected and classified for a given method. B) Proteins classified by SVM or TAGM-MAP were cross-referenced against the Human Protein Atlas and any protein considered to be singularly localized with an Enhanced rating was kept. The percentage of these proteins that were consistently classified by SVM or TAGM-MAP into the HPA-designated organelle is shown. Numbers above bars indicate the number of HPA proteins considered for each organelle. For conditions with Unknown proteins excluded, those proteins that were consistently classified as Unknown were removed from the analysis.

We did a similar analysis for additional organelles by comparing to the Human Protein Atlas (HPA)^25^. To obtain a baseline to our analysis, we focused on those proteins considered by the HPA to be localized to a single organelle with high confidence (enhanced rating). Of those proteins, we then plotted the percentage that were similarly classified by SVM or TAGM-MAP (Fig. 5B). Again, we found that TAGM-MAP outperformed SVM for mitochondrial proteins, and we saw a similar trend for ER proteins, albeit to a lesser degree. Conversely, SVM outperformed TAGM-MAP in the Golgi apparatus, nucleus, peroxisomes, and plasma membrane, although only two proteins were considered for the peroxisome. Similar to our observations above, the exclusion of unknown proteins yielded a larger increase in percentage agreement for SVM (orange vs blue bars) than TAGM-MAP (dark orange vs green bars); this exclusion also increased the performance in the cytosol for SVM over TAGM-MAP. These data correspond well to those of Figure 4A where 114 proteins designated as unknown by SVM were classified as mitochondrial by TAGM-MAP. Similar trends were found within ΔNef data (Fig. S8). Taken together, this suggests that at least in this cell system and using these fractionation methods, TAGM-MAP is better suited for spatial proteomic studies focused on the mitochondria and the ER, while SVM is better suited for studies of the Golgi, nucleus, and plasma membrane. This finding was surprising as we observed higher average QSep scores for the mitochondria and ER in WT replicates using SVM as compared to TAGM-MAP (Fig. S9), with less of a difference in ΔNef replicates (Fig. S10), which suggests an imperfect correlation between QSep scores and general accuracy for certain organelles.

### SVM-based BANDLE of WT replicates yielded the best agreement of HIV-dependent translocations with known HIV interactomes; partial overlap with ΔNef translocation hits

Following our analysis of classifiers, we examined various pipelines for identifying protein translocations. We inputted our SVM and TAGM-MAP classified data into BANDLE^10^ and a basic label-based analysis^7^, and inputted unclassified data into TRANSPIRE^9^ (Fig. 2A). For TRANSPIRE, we combined the organelles into 5 groups: 1) Golgi apparatus/plasma membrane/endoplasmic reticulum/peroxisomes/lysosomes, 2) cytosol/actin cytoskeleton/proteasome, 3) nucleus, 4) mitochondria, and 5) 40S/60S ribosome. This is in line with the authors’ recommendation to combine similarly behaving organelles to increase translocation confidence^9^, although in our case we lose the ability to identify proteins moving between the membranous organelles most likely to be affected by Nef, i.e. secretory organelles. To compare the performance of these five methods, we cross-referenced their hits against an HIV interactome derived from affinity purification-mass spectrometry (AP-MS)^26^ as well as the NIH HIV interactome^27^. The AP-MS study is more stringent since it includes only those proteins that directly complex with HIV proteins, while the NIH HIV interactome includes proteins that are affected by HIV even in the absence of evidence for a direct interaction. We found that the percentage of hits from each method that were in the interactomes was consistently above the threshold expected by chance (Fig. 6A, dashed line). Comparing the methods, the top 50 hits from the BANDLE analysis of SVM-classified data performed best for both interactomes with 20% and 84% of hits in the Jäger et al study (direct interactome by AP-MS) and NIH HIV interactome (functional as well as direct interactors), respectively. Of note, ∼1,500 proteins were considered to be translocation hits by the BANDLE analysis of SVM, i.e. greater than 95% probability of translocation. The validity of this value is difficult to gauge, but it is much higher than the ∼50 proteins from the BANDLE analysis of TAGM-MAP-classified data with a similar probability of translocation. We conducted a similar hit analysis on our ΔNef inducible line and found that SVM-based BANDLE was still the highest performer for the NIH HIV interactome but was only 3^rd^ best for the AP-MS interactome (Fig. S11).

**Figure 6:**
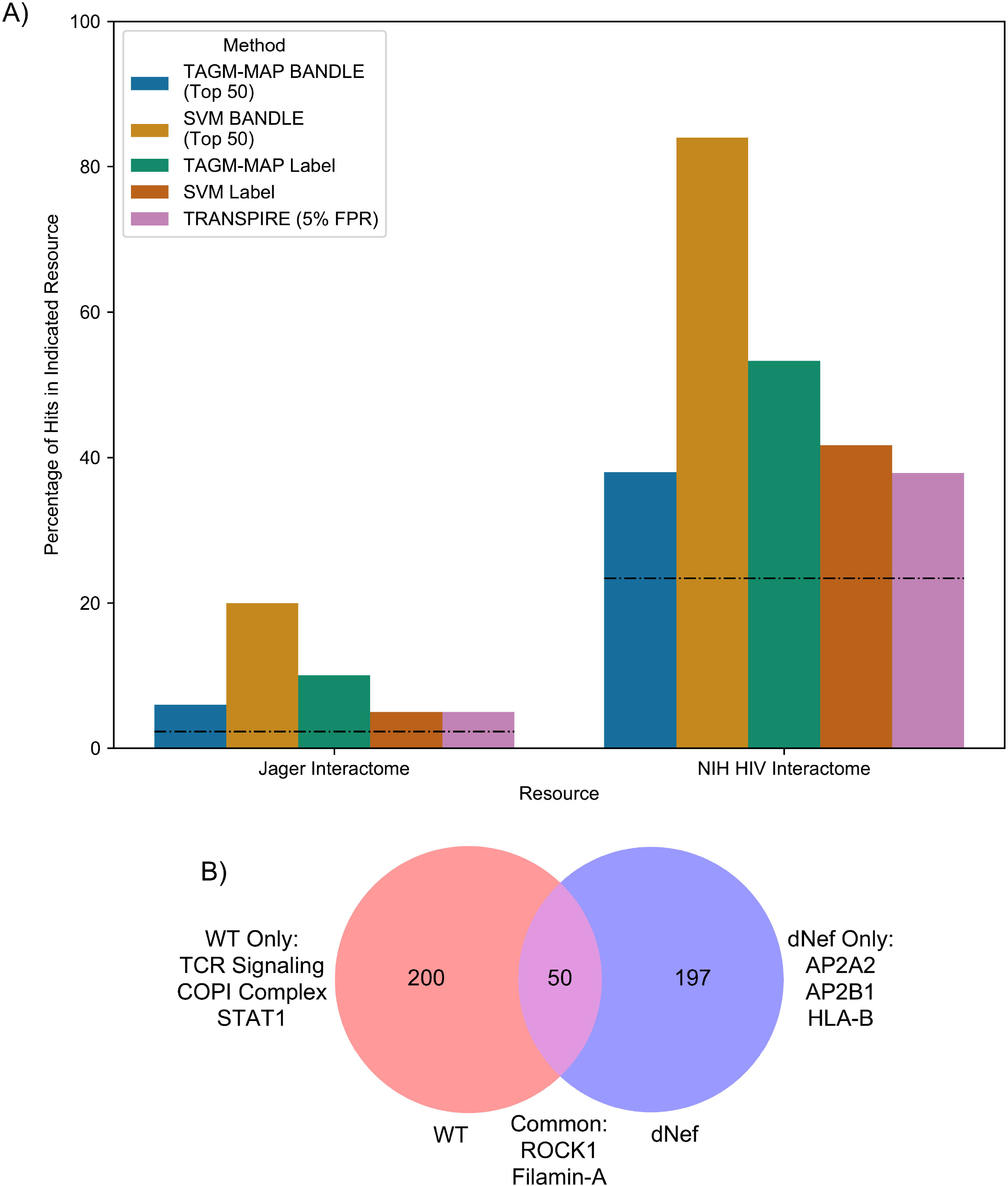
Detection of protein translocations by BANDLE analysis of SVM-classified data shows the highest rate of identifying known HIV interactors. A) The percentage of hits from each method that are in the Jäger HIV interactome^26^ (left bars) or the NIH HIV interactome^27^ (right bars) is shown. Dashed lines indicate the percentage of hits that would be expected by chance based on the proportion of the human proteome represented in each interactome. B) Venn diagram of top 250 hits from SVM-based BANDLE for WT and ΔNef replicates. Three of the hits from the ΔNef analysis were not detected by MS in WT replicates and were thus removed from consideration.

The top 250 hits from SVM-based BANDLE for WT and ΔNef were compared to see if the method could identify Nef-dependent translocations (Fig. 6B); hits that were detected by MS in only WT or ΔNef replicates were removed to avoid detection bias. Of those hits found only for WT, we observed several known Nef targets and cofactors: ZAP70 (ref.^37^), Lck^18, 38^, STAT1 (ref.^39^), and coatomer complex I (COPI complex)^40, 41^. Five separate proteins in the COPI complex appear together as well as three proteins from the T cell signaling pathway, suggesting high coverage of perturbed complexes. For commonly shared hits, proteins involved in cytoskeletal organization were enriched. Disruption of the cytoskeleton following infection with HIV has been attributed to Nef among other viral proteins, but the enriched proteins here lacked known targets of Nef but instead included ROCK1, an interactor of HIV Tat, and filamin-A, an interactor of HIV Gag^42^. We were surprised to see two components of the AP2 complex, known interactors of Nef^43^, and HLA class B, a known target of Nef^44, 45^, in the ΔNef only translocations. The SVM classification for these select proteins and STRING diagrams of the full protein sets are shown in the Supplemental Figures (Fig. S12-15). Notably, the SVM classifications rarely provided definitive organellar translocations for the hits identified by BANDLE (Fig. S12). In some cases, this was due to the majority of replicates becoming unclassified in the induced condition. An interesting exception is Filamin-A: although a translocation hit in both WT and ΔNef cells by BANDLE (Fig. 6B), by SVM classification Filamin-A moves from the actin cytoskeleton to the cytosol in cells expressing WT but not ΔNef (Fig. S12K). While the basis for such analytic discrepancies is unclear, taken together these data suggest potential value in identifying novel HIV cofactors, targets, and interactors via BANDLE analysis of spatial proteomics data.

## Discussion

We have detailed here a comparison of computational methods within the field of spatial proteomics as an example and guide for researchers hoping to use these methods to better understand viral infection and replication. Extensive work in the field, particularly by the Lilley^2, 11, 21, 36, 46^, Cristea^7–9^, and Borner groups^4–6, 47^ along with their collaborators, offers a variety of established choices for fractionation, classification, and translocation identification methods. To build off of the work of the Cristea group with HCMV, we chose to examine HIV-1 as a model virus due to the existing wealth of knowledge on HIV-dependent protein interactions and translocations. We found in our T cell line model and using differential centrifugation for cell fractionation that the choice of computational method for classification is organelle-dependent: TAGM-MAP offered an advantage for mitochondrial and ER proteins, while SVM performed better for the Golgi apparatus, nucleus, and plasma membrane. For identifying translocations, BANDLE gave the highest agreement with known HIV biology (i.e. published interactome data), particularly when coupled with SVM-classified data.

The model of inducible HIV in Jurkat T cells used here has advantages and disadvantages. One advantage is that the system provides a highly homogenous population of HIV-expressing cells suitable for mass spectrometric analysis^13^. A homogenous population is particularly important in spatial proteomic studies as mixed populations of cells might yield erroneous classifications of proteins due to mixtures of different states^46^. Another advantage is scalability. These experiments required just over 3×10^8^ cells for each technical replicate, or over 1×10^9^ cells for a single biological replicate, to be induced. In our initial attempts with fewer cells, centrifugation at higher RCF (110,000x*g* and 195,500x*g*) yielded insufficient protein mass for quality control and mass spectrometry (data not shown). This highlights an advantage of using this T cell line compared to using primary CD4+ T-cells^48^, which in principle would be more relevant but would require at least 2×10^9^ cells and extraordinary viral inocula to achieve a high-multiplicity, synchronized infection. A disadvantage of using this T cell system is that the cytoplasmic volume of the cell is relatively small. We required an order of magnitude more cells for each technical replicate here than was used in the D.O.M. studies of Itzhak et al., who used HeLa cells with larger cytoplasm.

In addition to these technical considerations for modeling viral infection/expression, the choice of fractionation method has practical and computational implications. The use of differential centrifugation here and by Itzhak et al. requires the downstream analysis of fewer fractions than gradient fractionation methods and is far less time-, resource-, and labor-intensive^2^. On the other hand, gradient fractionation methods seem to show increased resolution of protein classification^21^. In an attempt to increase the organellar resolution of the D.O.M method we used additional high-speed centrifugation steps to those described in the D.O.M. method of Itzhak et al. and found a significant increase in overall organellar resolution using seven fractions as compared to the original five (Fig. 2). Previous work by the Lilley group comparing differential centrifugation and gradient-based methods for fractionation revealed comparable downstream results for the two methods using U-2 OS cells with differential centrifugation having a slight advantage in resolving the cytosol and proteasome^36^, but whether this trend would hold in different cell types after viral infection or gene-expression is unclear. Generalizable rules for spatial proteomics might require comparisons of various fractionation and computational methods in multiple systems, or perhaps more likely, the specific experimental system and questions asked might be best addressed by a specific method. For example, to investigate translocations caused by HIV-1 Nef, better separation of membranous organelles (see Fig. S5 and S6) might have yielded more Nef-specific translocations.

Our findings on classification consistency and accuracy might influence the choice of classifier, at least for this model system. We found that SVM yielded higher consistency in classification than TAGM-MAP, although both suffered similar losses in consistency following HIV expression. In cases where infection or viral expression is expected to introduce greater noise in the data, as seems to be the case here, SVM may be the better option as it yielded a higher starting point for consistency. If lower tolerance to noise is acceptable, TAGM-MAP offers an advantageous alternative for both the mitochondria and ER. TAGM-MAP also suffered less loss of protein classification to unknown designations for uninduced replicates, perhaps due in part to the threshold used here for retaining SVM classification. While we used a basic median SVM algorithm score threshold for each organelle^2^ to allow for raw comparisons of classifiers to existing spatial proteomes, this might have been overly stringent for certain organelles, which would explain the higher number of unknown designated proteins for SVM. An alternative method would be to introduce an organelle-dependent threshold that would cap false positives by comparing classifier outputs to gene ontology analysis and published spatial proteomes; this method was employed previously by the Lilley group^1, 36^. We further note the fact that while SVM showed generally higher QSep scores for the mitochondria and ER it still underperformed compared to TAGM-MAP for these organelles. This suggested to us that organellar resolution as measured by QSep might be an imperfect measure of classification accuracy for a given organelle, a hypothesis that will need further examination.

Lastly, the choice of translocation identification method requires consideration of several factors, the first of which is the experimental design. Part of BANDLE’s power comes from its ability to factor multiple replicates of a condition into hit determination. Indeed, we saw a generally higher predictive power for BANDLE compared to other methods. The ranked list of output is also useful in cases where resources are limited and only a few hits can be pursued. TRANSPIRE seemed to have poorer performance compared to other methods, but this might reflect our need to combine similarly fractionated organelle groups to reduce computational demand and increase resolution. In cases where individual organellar resolution is greater, TRANSPIRE might yield higher quality hits. Notably, both BANDLE and TRANSPIRE require intensive computational resources, with TRANSPIRE requiring supercomputer access for larger, more complex datasets. In cases where computational power is limited, label-based methods would be suitable. Indeed, this method was employed by the Cristea group for their HCMV study with success^7^.

A challenge not addressed here is how to handle changes in whole-organellar behavior within spatial proteomics, such as might be induced by viruses. Indeed, we observed such a change within our data: peroxisomal marker proteins shifted in their fractionation behavior (peak abundance occurring at a higher *g*-force) when WT HIV was induced, becoming very similar in their behavior to marker proteins of the ER (Fig. S5). This effect was not observed for ΔNef (Fig. S6). A previous discussion of this issue by the Lilley group^10^ highlighted the various possible causes of whole-organellar changes—e.g. differences in organelle protein content, lipid composition, morphology, etc.—as potentially problematic for the movement-reproducibility method of translocation identification^5^, but how these types of biochemical changes would affect translocation detection methods or classifiers is not obvious. In our preliminary analyses of the average distance between organellar clusters based on pairwise distances, we found that peroxisomes alone shifted in relation to other organelles following the induction of WT HIV (but not ΔNef). However, analyses using QSep, which additionally considers the average intracluster distance (i.e., the dispersal of the cluster that defines the organelle), gave a less clear picture, with the potential for multiple relative movements among organelles (data not shown). These observations suggest that computational methodology will affect conclusions about organellar behavior as a whole. While the uniform shift of all markers for a given organelle should have only a minor impact on classification, how likely such a shift is in the context of viral gene expression probably depends on the specific virus and the type of cytopathic effect it induces. Indeed, the greater sensitivity of TAGM-MAP to HIV expression for classifier consistency could be a manifestation of subtle changes in organelle behavior. Careful examination of marker proteins used as well as the integration of pre-existing knowledge on the cytopathic effects of the virus under study are doubtlessly important for interpretation of whole-organellar changes.

With these considerations in mind, our findings underscore that studies of spatial proteomics require careful consideration of the question at hand to inform the choice of methodology. Our work and that of others highlights the potential differences in organellar resolution that can result from the choice of fractionation and analytical methods. Interest in a particular organelle and in specific types of translocations will factor into the choice of methods. Our findings offer a reference point for studies of viral infection by spatial proteomics, for general studies of the spatial proteome, and for the study of additional gene dropout mutants of HIV-1.

## Supporting information

Protein ID Tables

## Abbreviations

(ACN): acetonitrile
(BANDLE): Bayesian analysis of differential localization experiments
(D.O.M.): Dynamic Organellar Mapping
(FBS): fetal bovine serum
(FACS): fluorescence-assisted cell sorting
(FA): formic acid
(G.O.): gene ontology
(HPA): Human Protein Atlas
(IAA): iodoacetamide
(MS): mass spectrometry
(PBS-T): PBS+0.1% Tween-20
(pen/strep): penicillin/streptomycin
(PCA): Principal Component Analysis
(SILAC): stable isotope labeling by amino acids in cell culture
(SVM): support vector machine
(TMT): tandem mass tag
(TAGM-MAP): t-augmented Gaussian mixture modeling with *maximum a posteriori* estimates
(TRANSPIRE): translocation analysis of spatial proteomics
(TCEP): tris(2-carboxyethyl)phosphine
(UPLC): ultra-performance liquid chromatography

## Data availability

Mass spectrometry data (.RAW files and peptide identification tables) can be found on the ProteomeXchange database using project accession number PXD024716.

## Supplemental data

This article contains supplemental data.

## Acknowledgements

The authors would like to thank the NIH AIDS Reagent Program and The Pendleton Charitable Trust. The authors would like to also thank Hannah Carter, Max Qian, and Oliver Crook for productive discussions on the computational analysis. This work was partially supported by a UC San Diego Center for AIDS Research (UCSD CFAR) Developmental Award, an NIH funded program (P30AI036214). A.LO. is supported by NIAID F31AI141111. C.A.S. is supported by a UCSD CFAR Developmental Award (P30AI036214). M.K.L. was supported by K08AI112394. A.R. and N.J.K. are funded by a grant to the HARC Center from the National Institutes of Health (P50AI150476 to N.J.K.). J.M.W. is supported by NIH/NIGMS Grants T32 GM007752 and NIH/NIAMS T32 AR064194. D.G. is supported by The Collaborative Center of Multiplexed Proteomics at UCSD. J.G. is supported by NIAID R01AI129706. All funding sources had no involvement in the design and execution of this study nor the final manuscript.

## Author contributions

A.L.O., C.A.S., M.K.L., and J.G. conceived the design and scope of this study. M.K.L. created the inducible cell lines used in this study. A.L.O. and C.A.S. performed the fractionation experiments. A.L.O., J.M.W., and D.G. performed pilot studies for the mass spectrometry analysis. A.R., K.S., and N.J.K. performed the mass spectrometry analysis. A.L.O. performed the computational analysis. A.L.O. and J.G. wrote the manuscript and all authors reviewed and edited the manuscript.

