## Supplemental Figures 1-15 for "Comparative Analysis of T Cell Spatial Proteomics and the Influence of HIV Expression"

Includes Supplemental Figures 1-15 (Protein ID Tables are in a separate .zip folder)

Figure S1

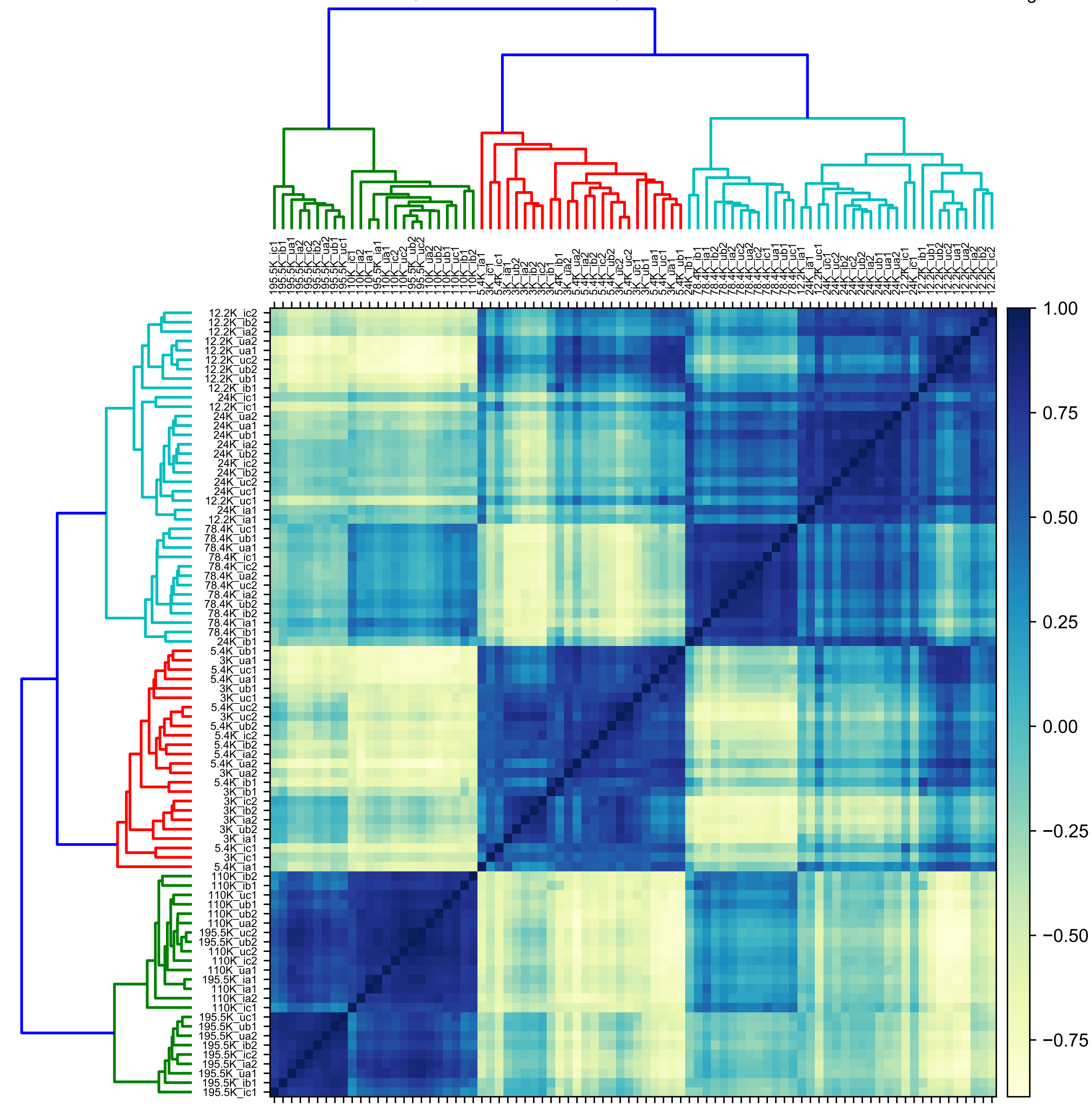

**Supplemental Figure S1:** Correlation analysis for WT cell fractions. Spearman correlation between individual fractions of both WT biological replicates. Fractions are labeled with a three-character code after the *g*-force: “u” or “i” for uninduced or induced, respectively; “a”, “b”, or “c” for the technical replicate; and “1” or “2” for the biological replicate. Only proteins detected in all WT replicates are considered here (4,765 proteins).

T Cell Spatial Proteomics and Impact of HIV

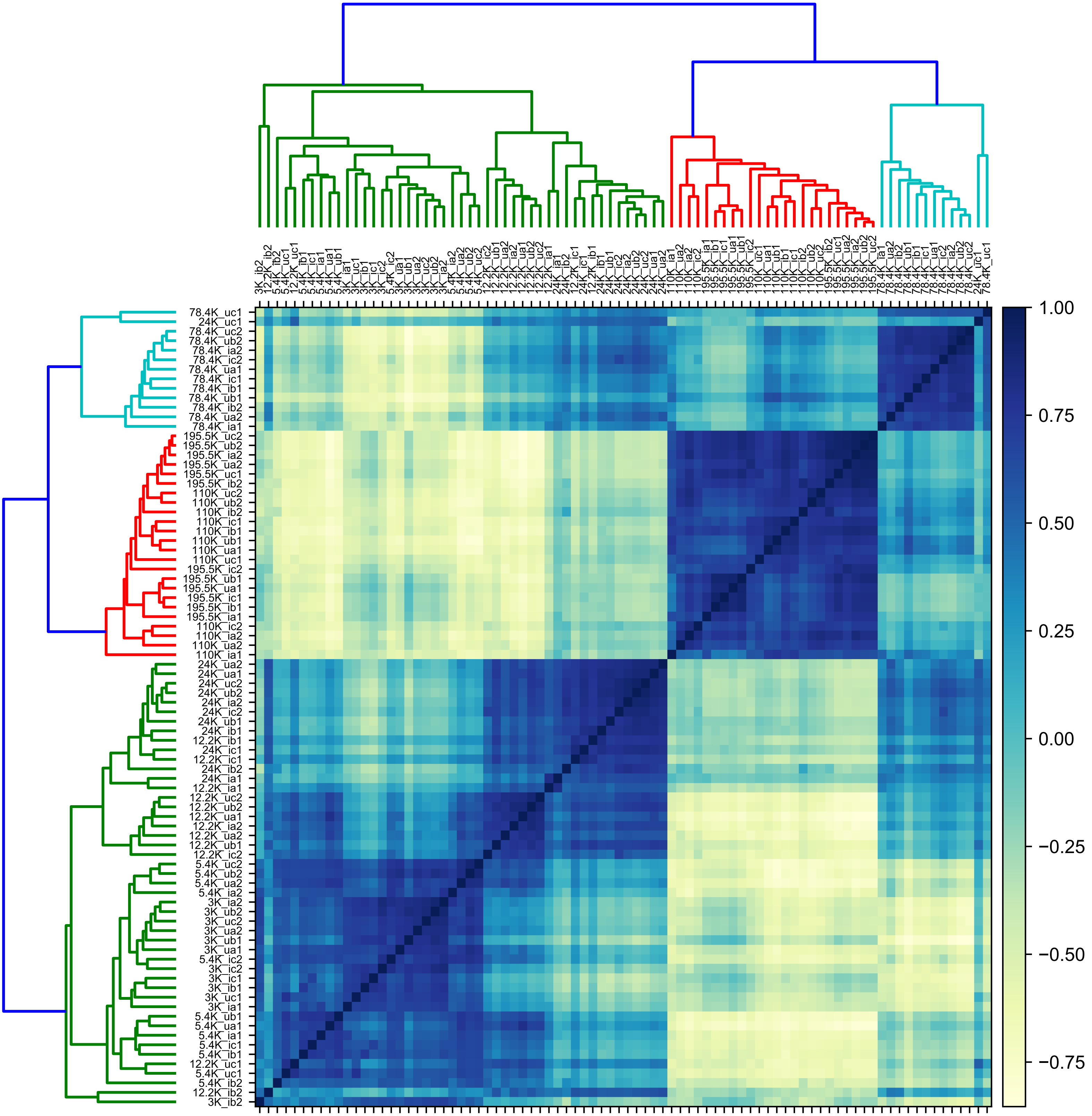

**Supplemental Figure S2:** Correlation analysis for ΔNef cell fractions. Spearman correlation between individual fractions of both ΔNef biological replicates. Fractions are labeled with a three-character code after the *g*-force: “u” or “i” for uninduced or induced, respectively; “a”, “b”, or “c” for the technical replicate; and “1” or “2” for the biological replicate. Only proteins detected in all ΔNef replicates are considered here (4,173 proteins).

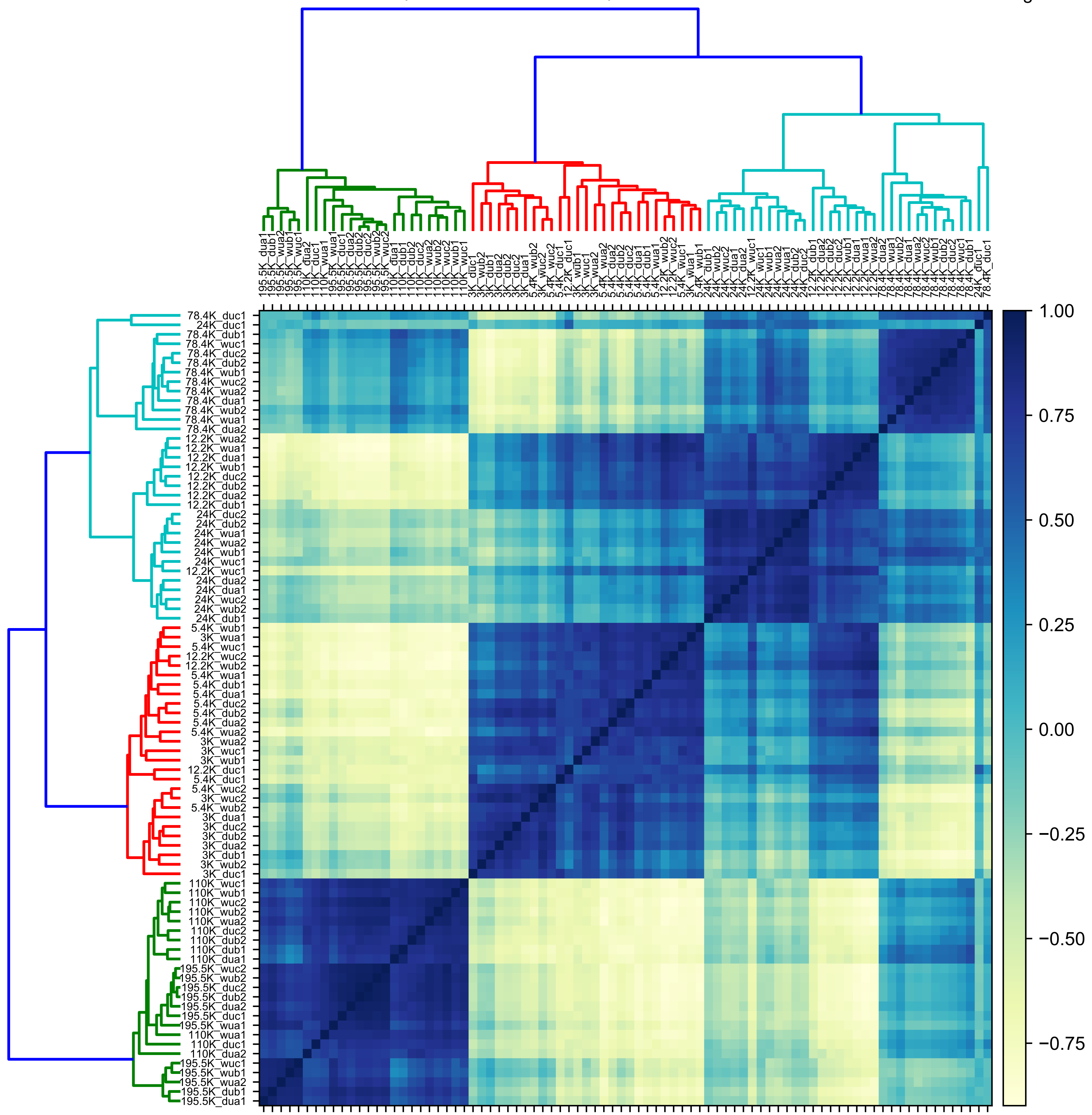
1.00
0.75
0.50
0.25
0.00
−0.25
−0.50
−0.75

**Supplemental Figure S3:** Correlation analysis for WT and ΔNef cell fractions. Spearman correlation between all individual fractions of uninduced condition. Fractions are labeled with a four-character code after the *g*-force: “w” or “d” for WT or ΔNef, “u” or “i” for uninduced or induced, respectively; “a”, “b”, or “c” for the technical replicate; and “1” or “2” for the biological replicate. Only proteins detected in all WT and ΔNef replicates are considered here (3,739 proteins).

Figure S4

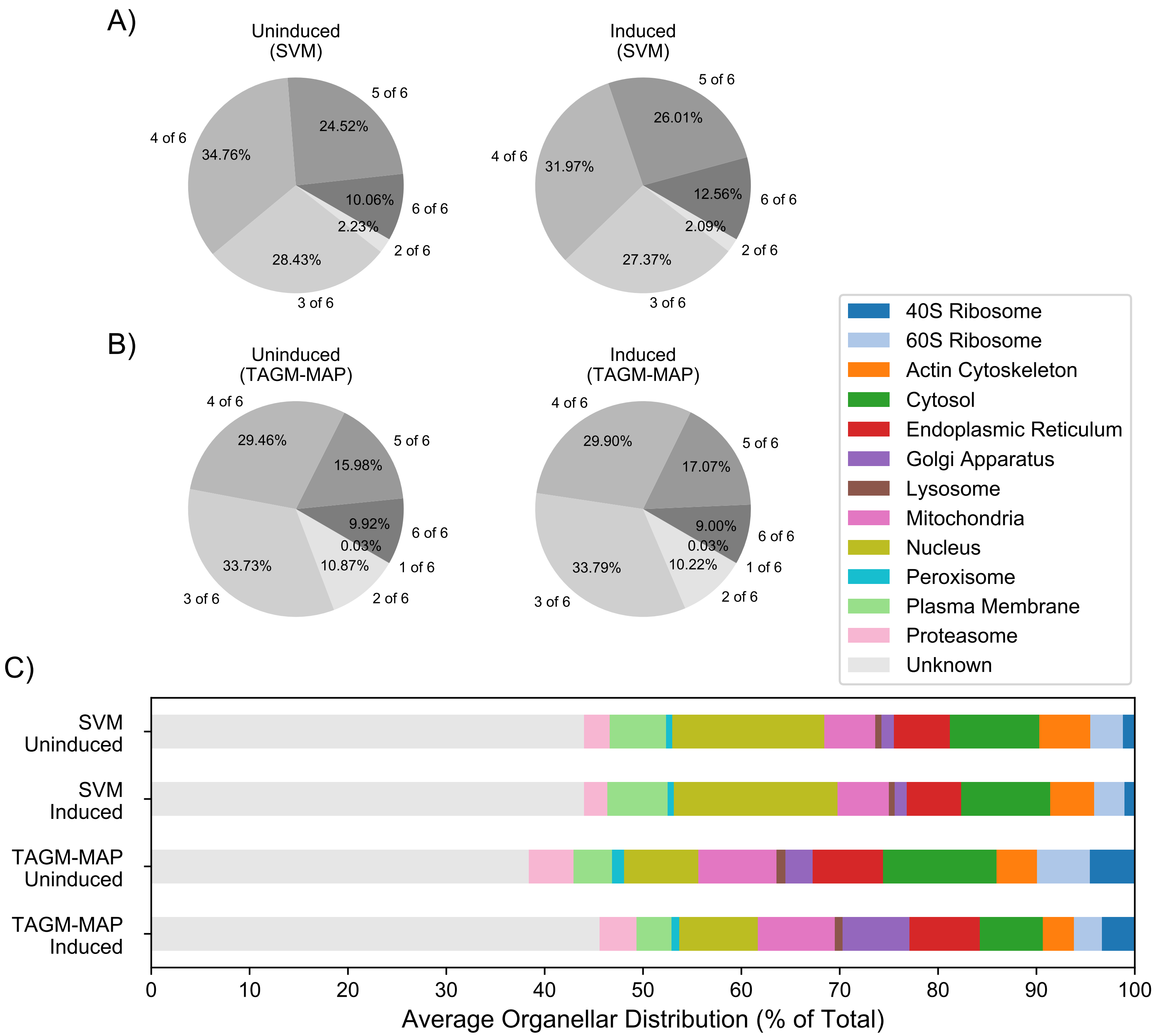
A)
Uninduced
(SVM)
5 of 6
24.52%
4 of 6
34.76%
10.06%
6 of 6
2.23%
2 of 6
28.43%
3 of 6
Induced
(SVM)
5 of 6
26.01%
4 of 6
31.97%
12.56%
6 of 6
2.09%
2 of 6
27.37%
3 of 6
B)
Uninduced
(TAGM-MAP)
4 of 6
29.46%
5 of 6
15.98%
9.92%
6 of 6
0.03%
1 of 6
33.73%
10.87%
3 of 6
2 of 6
Induced
(TAGM-MAP)
4 of 6
29.90%
5 of 6
17.07%
9.00%
6 of 6
0.03%
1 of 6
33.79%
10.22%
3 of 6
2 of 6
C)
40S Ribosome
60S Ribosome
Actin Cytoskeleton
Cytosol
Endoplasmic Reticulum
Golgi Apparatus
Lysosome
Mitochondria
Nucleus
Peroxisome
Plasma Membrane
Proteasome
Unknown
SVM Uninduced
SVM Induced
TAGM-MAP Uninduced
TAGM-MAP Induced
0
10
20
30
40
50
60
70
80
90
100
Average Organellar Distribution (% of Total)

**Supplemental Figure S4:** Classification with SVM shows greater consistency than TAGM-MAP classification for ΔNef replicates. A) Proteins were classified by SVM and the most frequent organellar classification was identified along with its frequency, i.e. number of technical replicates classified as such. Left pie chart shows consistency of classification for WT uninduced replicates and right pie chart shows WT induced replicates. B) Same as A), but classification by TAGM-MAP. C) Average distribution of proteins across organelles for each indicated condition. All charts consider the same common proteins found across all ΔNef replicates (4,173 proteins).

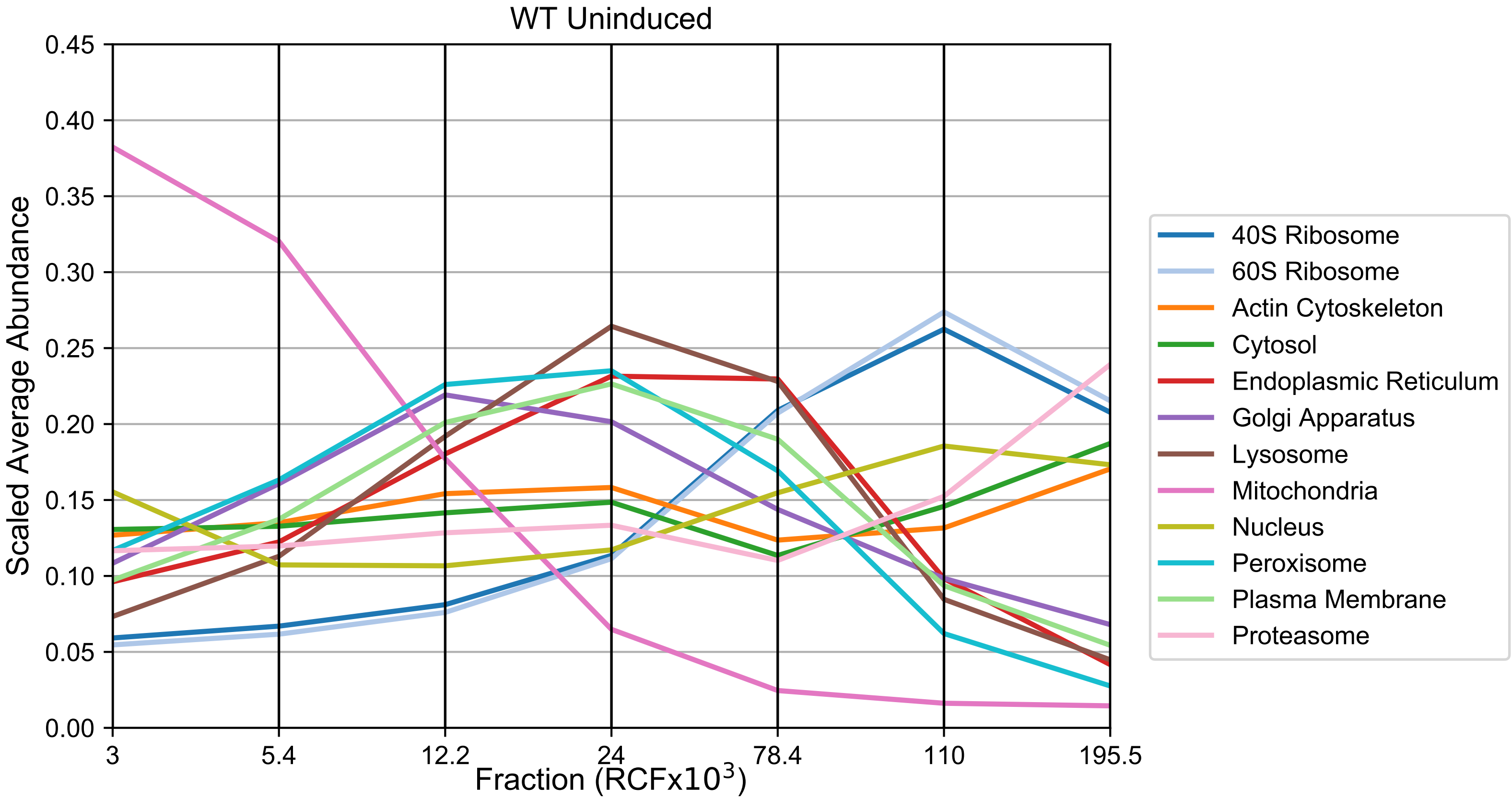
WT Uninduced
Scaled Average Abundance
0.45
0.40
0.35
0.30
0.25
0.20
0.15
0.10
0.05
0.00
3
5.4
12.2
24
78.4
110
195.5
Fraction (RCFx10$^3$)
40S Ribosome
60S Ribosome
Actin Cytoskeleton
Cytosol
Endoplasmic Reticulum
Golgi Apparatus
Lysosome
Mitochondria
Nucleus
Peroxisome
Plasma Membrane
Proteasome

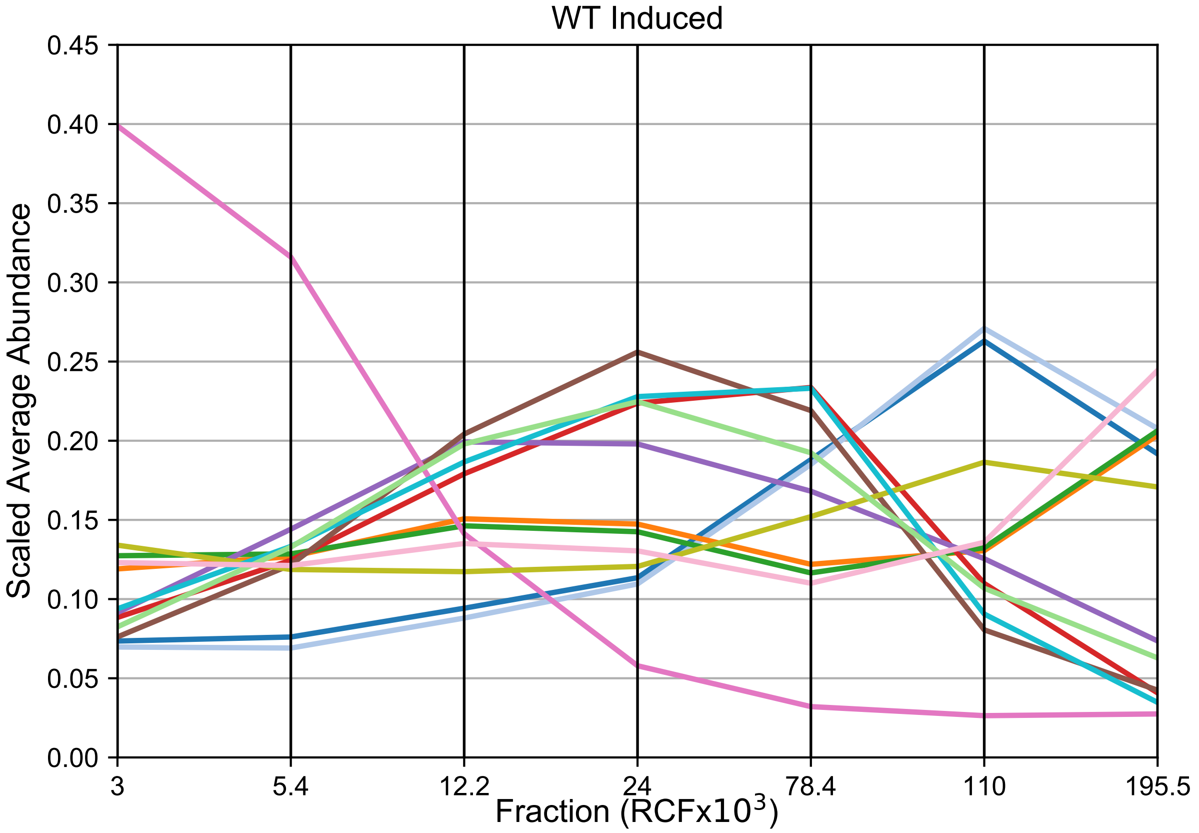
WT Induced
Scaled Average Abundance
0.45
0.40
0.35
0.30
0.25
0.20
0.15
0.10
0.05
0.00
3
5.4
12.2
24
78.4
110
195.5
Fraction (RCFx10$^3$)

**Supplemental Figure S5:** Marker profiles of WT replicates. Average behavior of markers for each organelle have been charted for uninduced (top) and induced (bottom) replicates. Values were row-normalized prior to averaging. The “hsap” marker set from pRoloc was used.

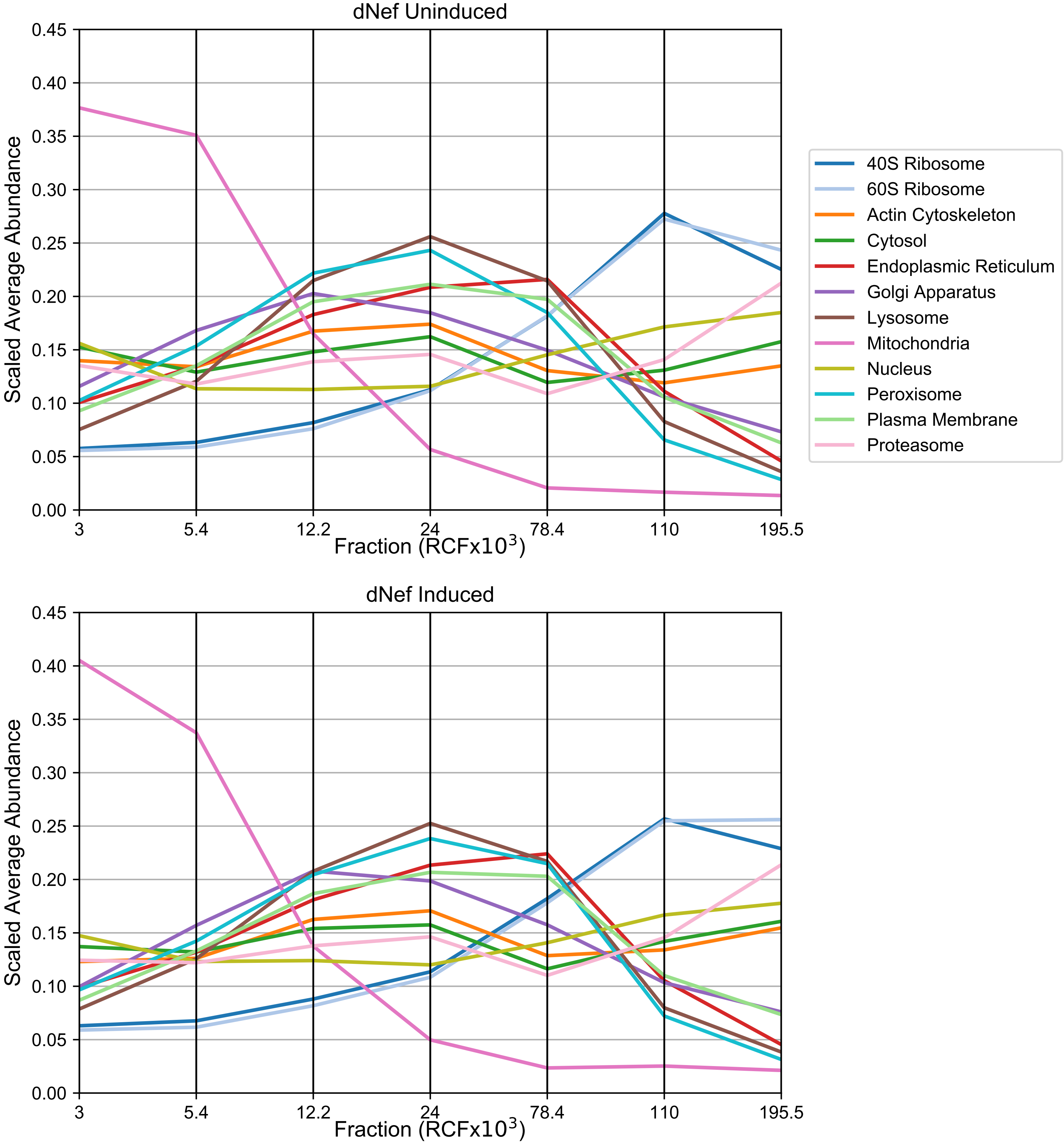
dNef Uninduced
Scaled Average Abundance
0.45
0.40
0.35
0.30
0.25
0.20
0.15
0.10
0.05
0.00
3
5.4
12.2
24
78.4
110
195.5
Fraction (RCFx10$^3$)
40S Ribosome
60S Ribosome
Actin Cytoskeleton
Cytosol
Endoplasmic Reticulum
Golgi Apparatus
Lysosome
Mitochondria
Nucleus
Peroxisome
Plasma Membrane
Proteasome
dNef Induced
Scaled Average Abundance
0.45
0.40
0.35
0.30
0.25
0.20
0.15
0.10
0.05
0.00
3
5.4
12.2
24
78.4
110
195.5
Fraction (RCFx10$^3$)

**Supplemental Figure S6:** Marker profiles of ΔNef replicates. Average behavior of markers for each organelle have been charted for uninduced (top) and induced (bottom) replicates. Values were row-normalized prior to averaging. The "hsap" marker set from pRoloc was used.

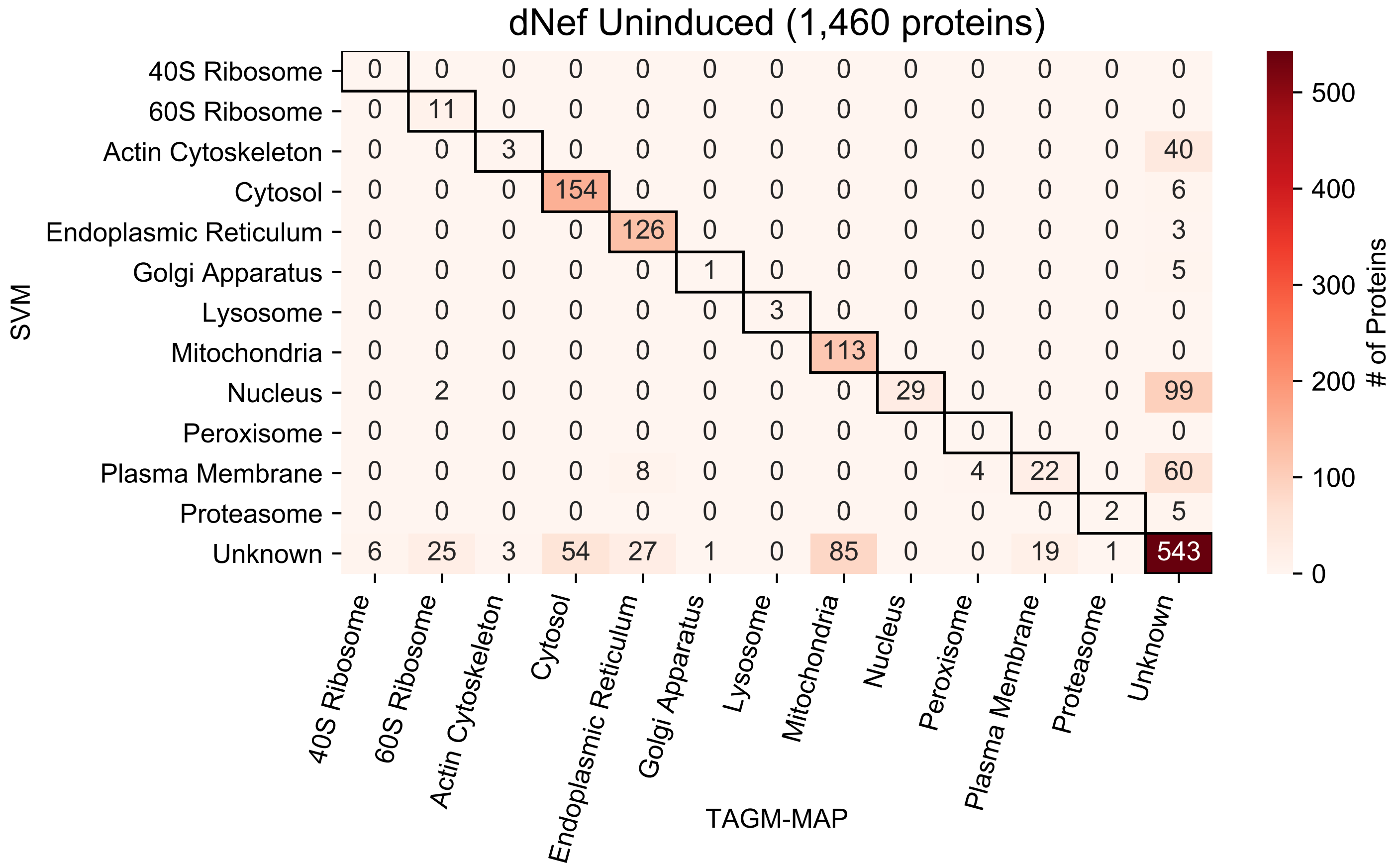
dNef Uninduced (1,460 proteins)
SVM
TAGM-MAP
### of Proteins
0
100
200
300
400
500
40S Ribosome 0 0 0 0 0 0 0 0 0 0 0 0 0
60S Ribosome 0 11 0 0 0 0 0 0 0 0 0 0 0
Actin Cytoskeleton 0 0 3 0 0 0 0 0 0 0 0 0 40
Cytosol 0 0 0 154 0 0 0 0 0 0 0 0 6
Endoplasmic Reticulum 0 0 0 0 126 0 0 0 0 0 0 0 3
Golgi Apparatus 0 0 0 0 0 1 0 0 0 0 0 0 5
Lysosome 0 0 0 0 0 0 3 0 0 0 0 0 0
Mitochondria 0 0 0 0 0 0 0 113 0 0 0 0 0
Nucleus 0 2 0 0 0 0 0 0 29 0 0 0 99
Peroxisome 0 0 0 0 0 0 0 0 0 0 0 0 0
Plasma Membrane 0 0 0 0 8 0 0 0 0 4 22 0 60
Proteasome 0 0 0 0 0 0 0 0 0 0 0 2 5
Unknown 6 25 3 54 27 1 0 85 0 0 19 1 543
40S Ribosome
60S Ribosome
Actin Cytoskeleton
Cytosol
Endoplasmic Reticulum
Golgi Apparatus
Lysosome
Mitochondria
Nucleus
Peroxisome
Plasma Membrane
Proteasome
Unknown

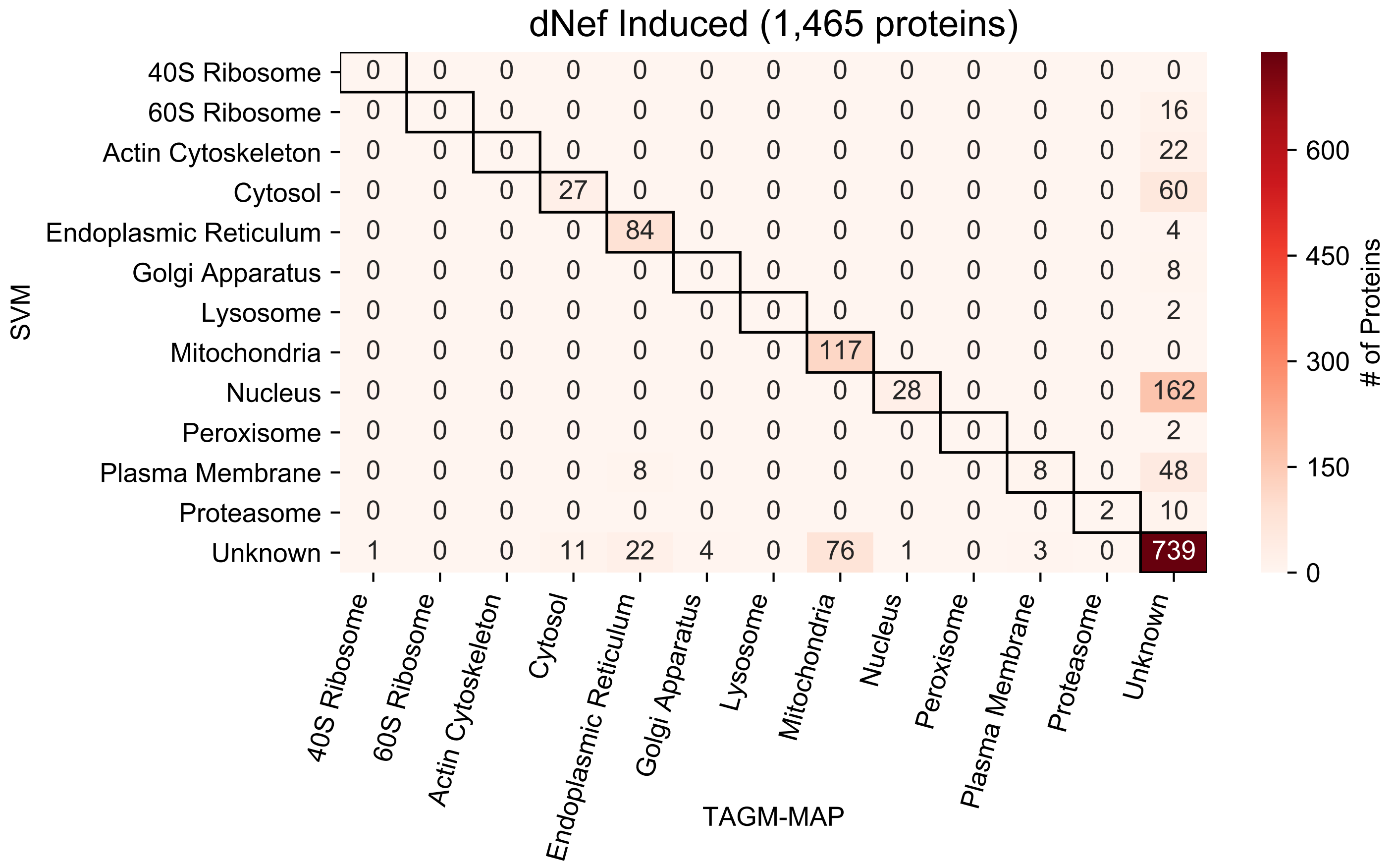
dNef Induced (1,465 proteins)
SVM
TAGM-MAP
### of Proteins
0
150
300
450
600
40S Ribosome 0 0 0 0 0 0 0 0 0 0 0 0 0
60S Ribosome 0 0 0 0 0 0 0 0 0 0 0 0 16
Actin Cytoskeleton 0 0 0 0 0 0 0 0 0 0 0 0 22
Cytosol 0 0 0 27 0 0 0 0 0 0 0 0 60
Endoplasmic Reticulum 0 0 0 0 84 0 0 0 0 0 0 0 4
Golgi Apparatus 0 0 0 0 0 0 0 0 0 0 0 0 8
Lysosome 0 0 0 0 0 0 0 0 0 0 0 0 2
Mitochondria 0 0 0 0 0 0 0 117 0 0 0 0 0
Nucleus 0 0 0 0 0 0 0 0 28 0 0 0 162
Peroxisome 0 0 0 0 0 0 0 0 0 0 0 0 2
Plasma Membrane 0 0 0 0 8 0 0 0 0 0 8 0 48
Proteasome 0 0 0 0 0 0 0 0 0 0 0 2 10
Unknown 1 0 0 11 22 4 0 76 1 0 3 0 739
40S Ribosome
60S Ribosome
Actin Cytoskeleton
Cytosol
Endoplasmic Reticulum
Golgi Apparatus
Lysosome
Mitochondria
Nucleus
Peroxisome
Plasma Membrane
Proteasome
Unknown

**Supplemental Figure S7:** Concordance of SVM and TAGM-MAP classifications for ΔNef replicates depend on organelle and expression of HIV. A) Heat map of common proteins that were consistently classified (proteins classified consistently in at least 4 of 6 replicates) by both SVM and TAGM-MAP for uninduced condition. Annotations indicate number of proteins in a given scenario. B) Same as A) for induced condition.

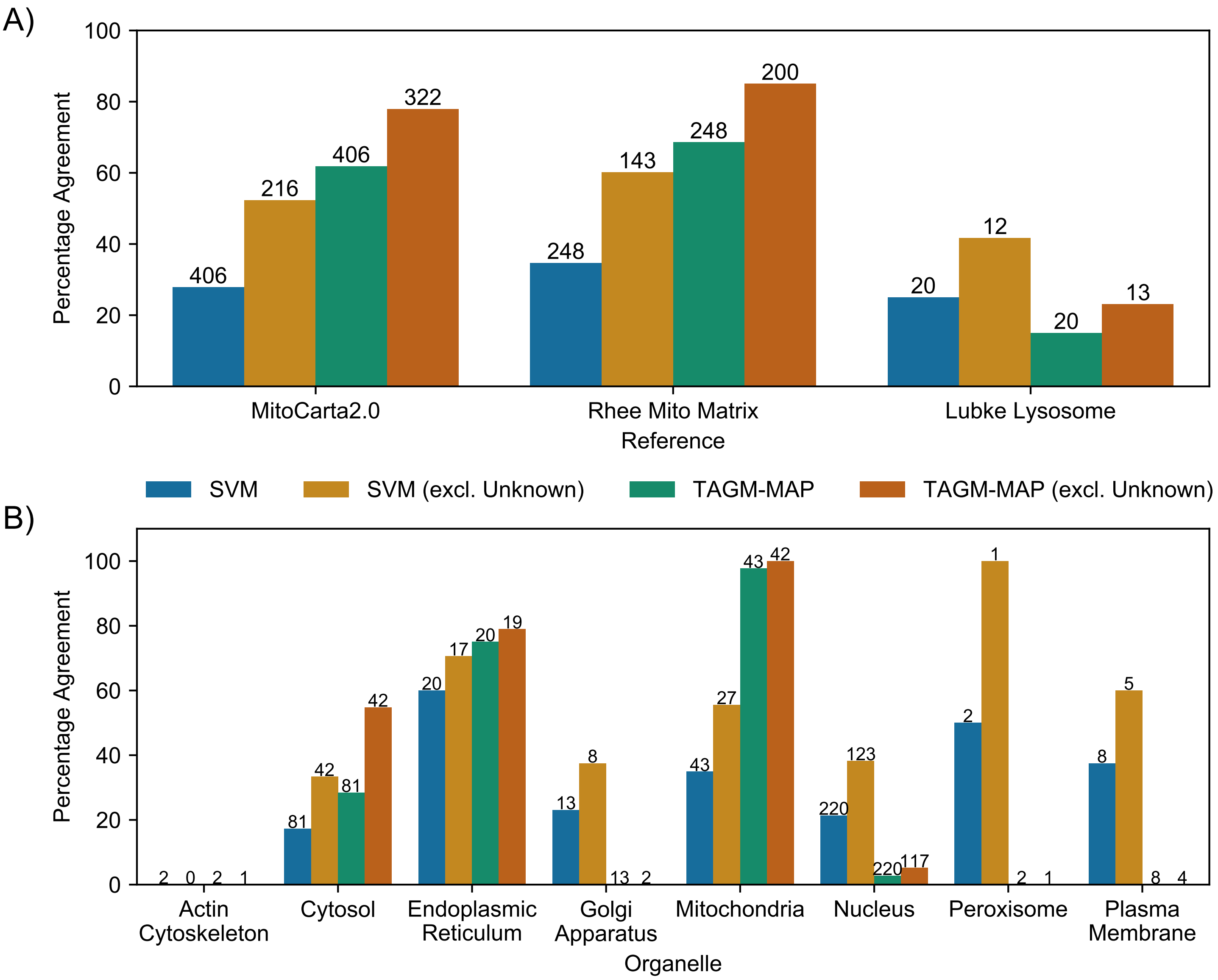
A)
Percentage Agreement
100
80
60
40
20
0
406
216
406
322
248
143
248
200
20
12
20
13
MitoCarta2.0
Rhee Mito Matrix
Lubke Lysosome
Reference
SVM
SVM (excl. Unknown)
TAGM-MAP
TAGM-MAP (excl. Unknown)
B)
Percentage Agreement
100
80
60
40
20
0
2
0
2
1
81
42
81
42
20
17
20
19
13
8
13
2
43
27
43
42
220
123
220
117
2
1
2
1
8
5
8
4
Actin Cytoskeleton
Cytosol
Endoplasmic Reticulum
Golgi Apparatus
Mitochondria
Nucleus
Peroxisome
Plasma Membrane
Organelle

**Supplemental Figure S8:** Validation of protein classification of ΔNef replicates reveals higher performance for ER and mitochondria using TAGM-MAP, but better performance for Golgi apparatus, nucleus, and plasma membrane using SVM. A) Percentage of detected proteins from MitoCarta2.0 database[22], Rhee et al. mitochondrial matrix study[23], or Lubke lysosome proteome[24] that were consistently classified (proteins classified consistently in at least 4 of 6 replicates) in line with the respective reference. Numbers above bars indicate the total number of proteins from that reference that were detected and classified for a given method. B) Proteins classified here were cross-referenced against the Human Protein Atlas and any protein considered to be singularly localized with an Enhanced rating was kept. The percentage of these proteins that were consistently classified by SVM or TAGM-MAP into the HPA-designated organelle is shown. Numbers above bars indicate the number of HPA proteins considered for each organelle. For conditions with Unknown proteins excluded, those proteins that were consistently classified as Unknown were removed from the analysis.

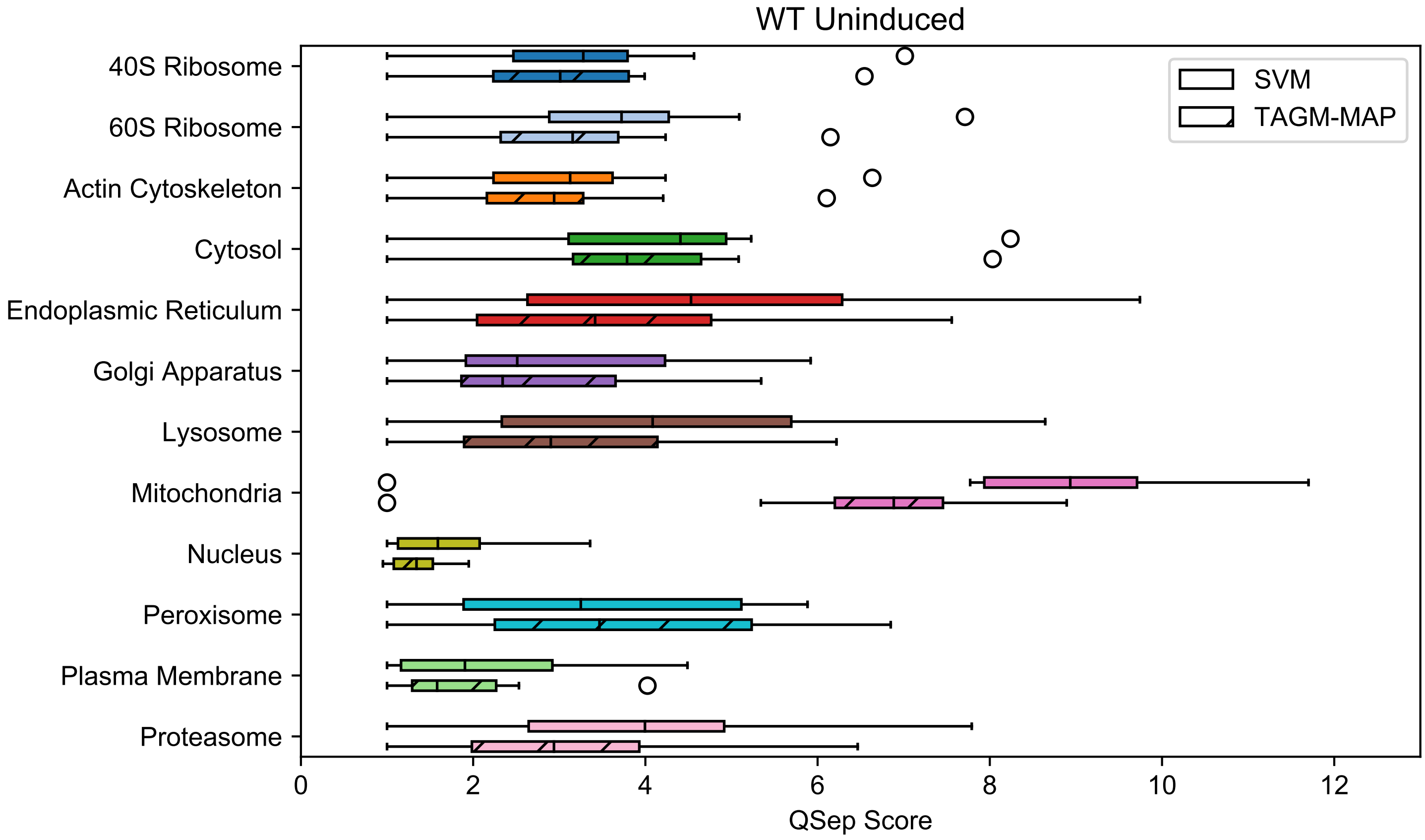
WT Uninduced
40S Ribosome
60S Ribosome
Actin Cytoskeleton
Cytosol
Endoplasmic Reticulum
Golgi Apparatus
Lysosome
Mitochondria
Nucleus
Peroxisome
Plasma Membrane
Proteasome
SVM
TAGM-MAP
0
2
4
6
8
10
12
QSep Score

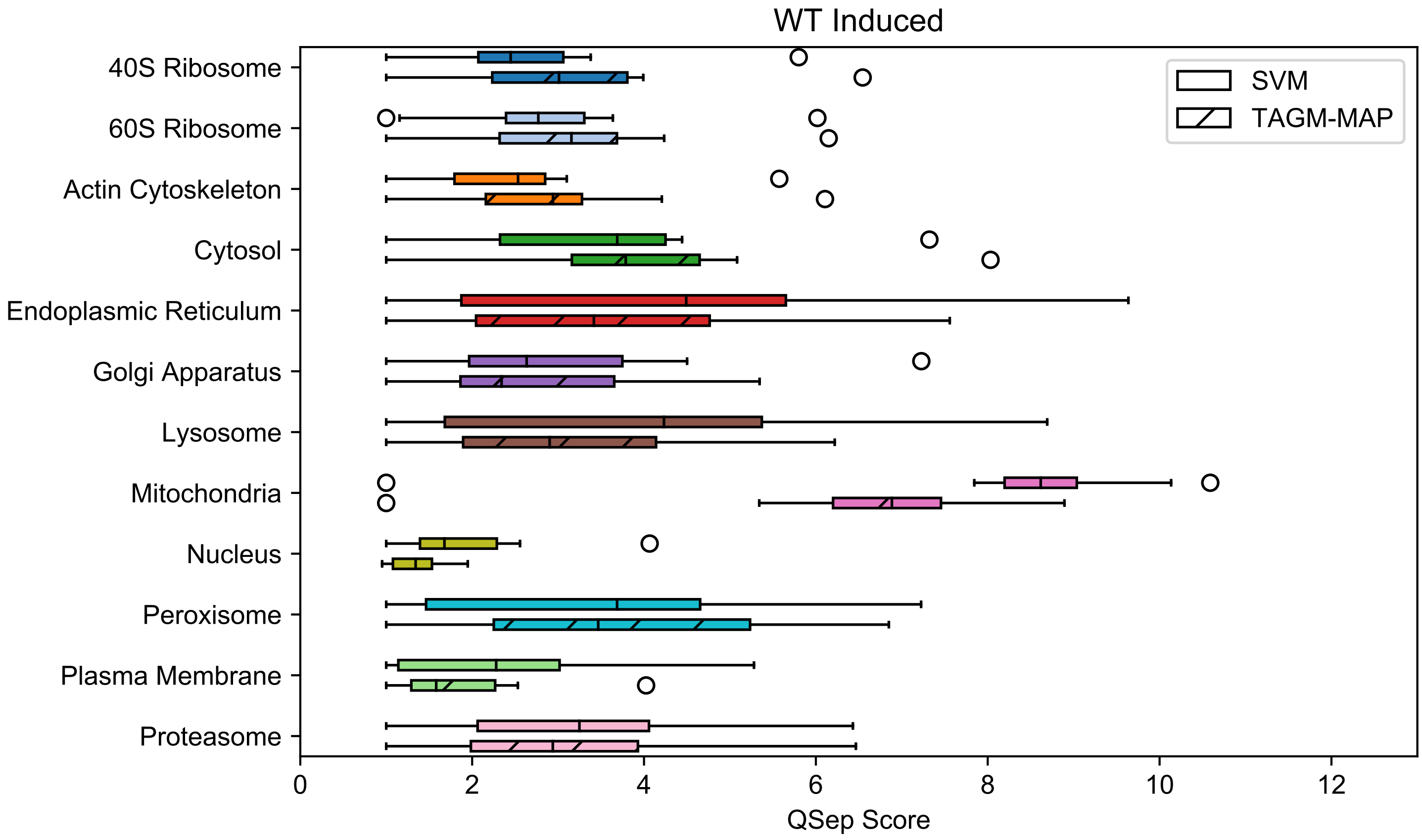
WT Induced
40S Ribosome
60S Ribosome
Actin Cytoskeleton
Cytosol
Endoplasmic Reticulum
Golgi Apparatus
Lysosome
Mitochondria
Nucleus
Peroxisome
Plasma Membrane
Proteasome
SVM
TAGM-MAP
0
2
4
6
8
10
12
QSep Score

**Supplemental Figure S9:** SVM classification shows generally higher organellar QSep scores for WT replicates. The individual organellar QSep scores, i.e. distance between organelle A and B, were averaged across uninduced replicates (top) or induced replicates (bottom) then plotted.

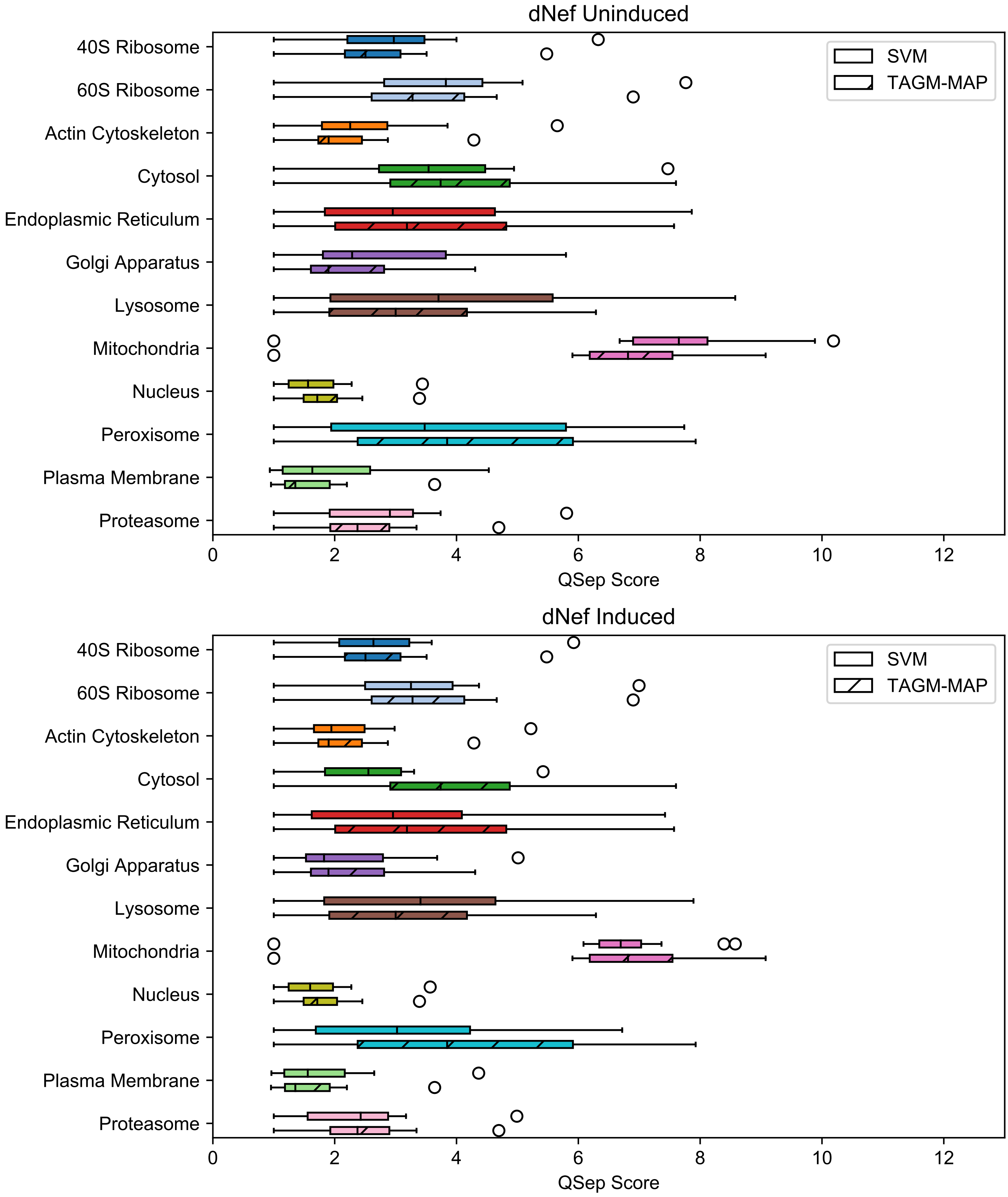
dNef Uninduced
40S Ribosome
60S Ribosome
Actin Cytoskeleton
Cytosol
Endoplasmic Reticulum
Golgi Apparatus
Lysosome
Mitochondria
Nucleus
Peroxisome
Plasma Membrane
Proteasome
SVM
TAGM-MAP
0
2
4
6
8
10
12
QSep Score
dNef Induced
40S Ribosome
60S Ribosome
Actin Cytoskeleton
Cytosol
Endoplasmic Reticulum
Golgi Apparatus
Lysosome
Mitochondria
Nucleus
Peroxisome
Plasma Membrane
Proteasome
SVM
TAGM-MAP
0
2
4
6
8
10
12
QSep Score

**<u>Supplemental Figure S10:</u>** SVM and TAGM-MAP classification show mixed results for organellar resolution by QSep scores for ΔNef replicates. The individual organellar QSep scores, i.e. distance between organelle A and B, were averaged across uninduced replicates (top) or induced replicates (bottom) then plotted.

Figure S11

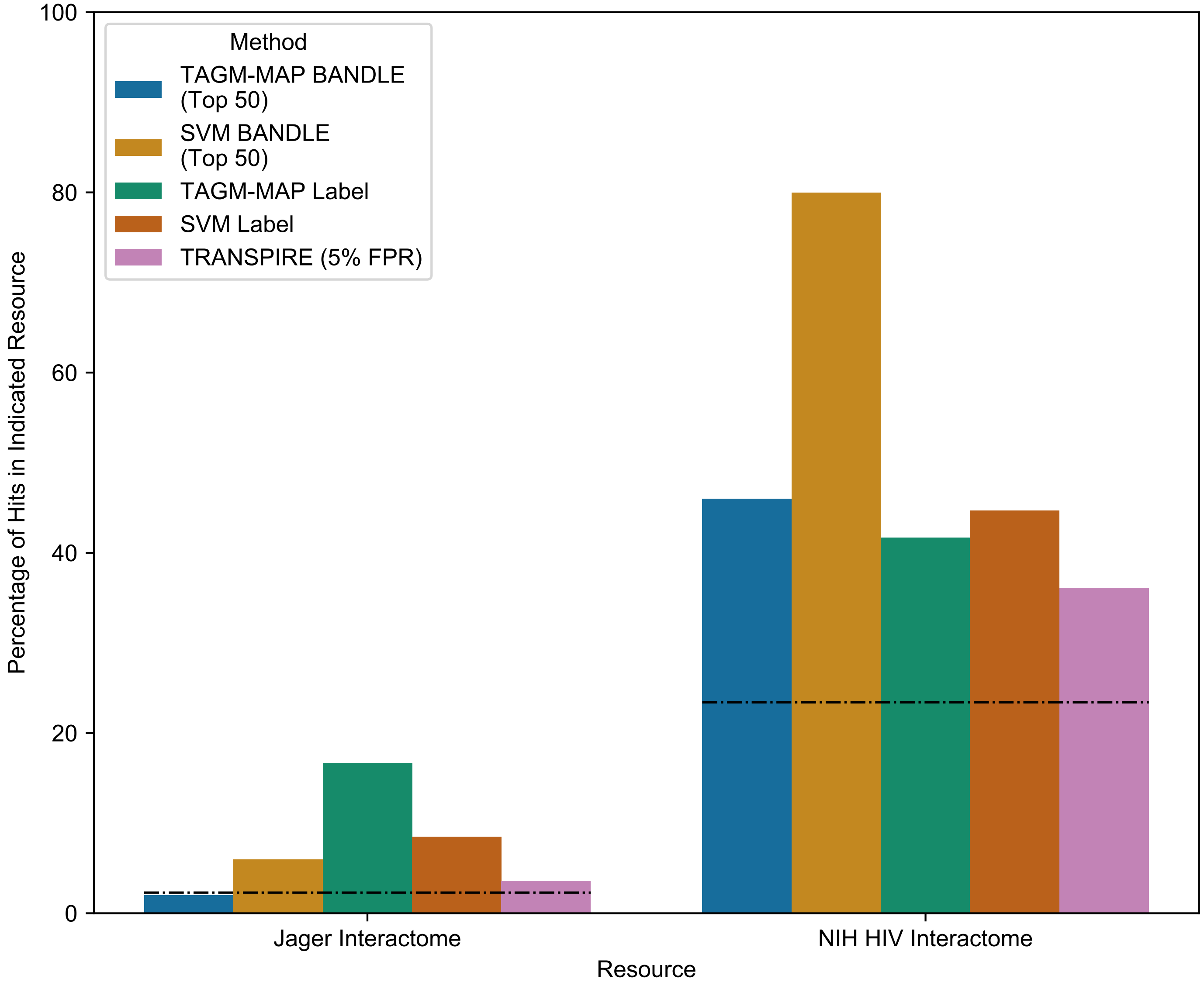
Method
TAGM-MAP BANDLE (Top 50)
SVM BANDLE (Top 50)
TAGM-MAP Label
SVM Label
TRANSPIRE (5% FPR)
Percentage of Hits in Indicated Resource
0
20
40
60
80
100
Jager Interactome
NIH HIV Interactome
Resource

**Supplemental Figure S11:** Performance of translocation identification method depends on the choice of known HIV interactors in ΔNef replicates. The percentage of hits from each method that are in the Jäger HIV interactome[26] (left bars) or the NIH HIV interactome[27] (right bars) is shown. Dashed lines indicate the percentage of hits that would be expected by chance based on the proportion of the human proteome represented in each interactome.

T Cell Spatial Proteomics and Impact of HIV

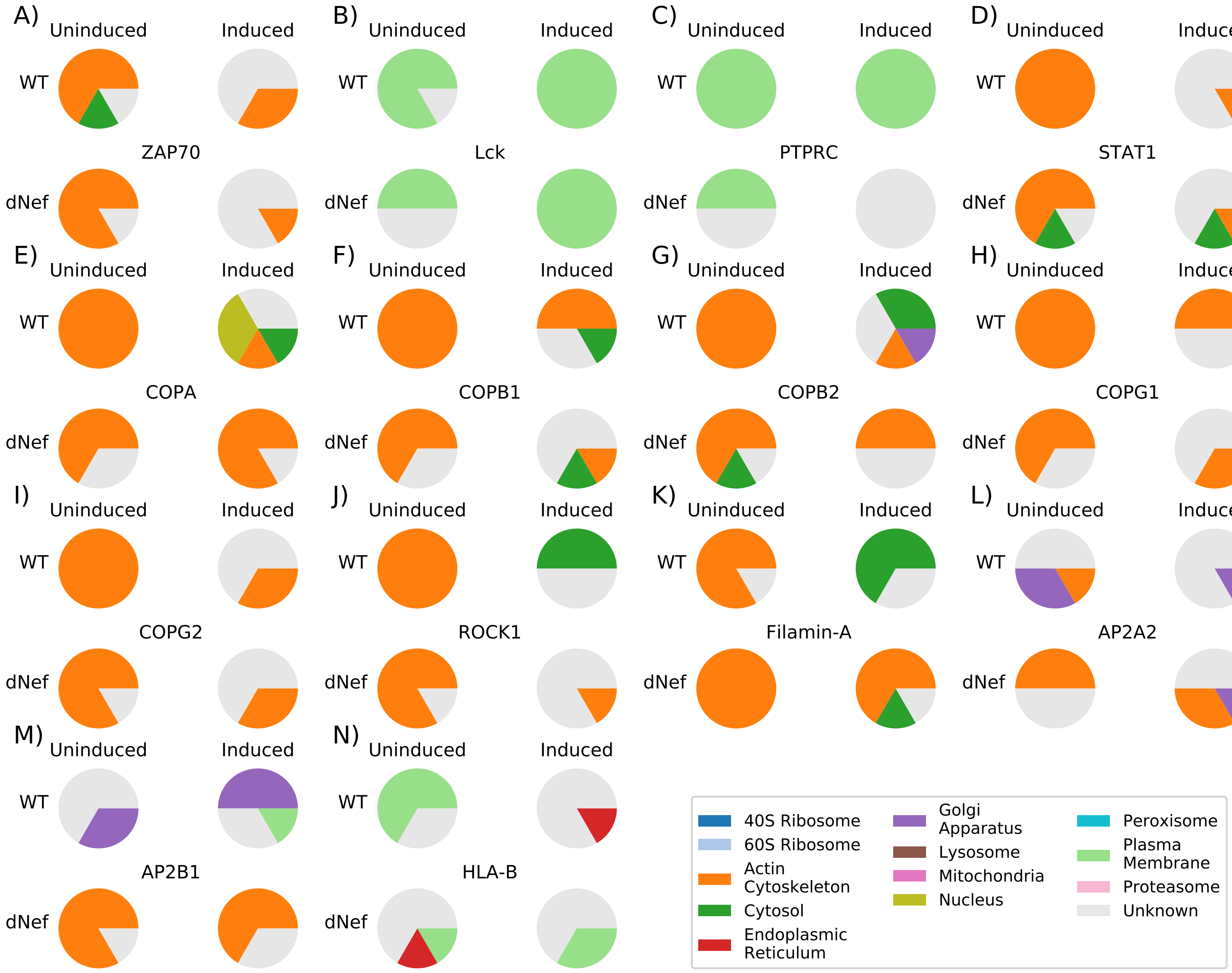

**Supplemental Figure S12:** SVM classification of selected SVM-based BANDLE hits. Pie charts displaying the classification across WT and ΔNef replicates for the WT only hits: TCR signaling proteins (A-C), STAT1 (D), and COPI complex (E-I). Cytoskeletal organizers ROCK1 (J) and Filamin-A (K) are common to both WT and ΔNef replicates. AP2A2 (L), AP2B1 (M), and HLA-B (N) were ΔNef only hits.

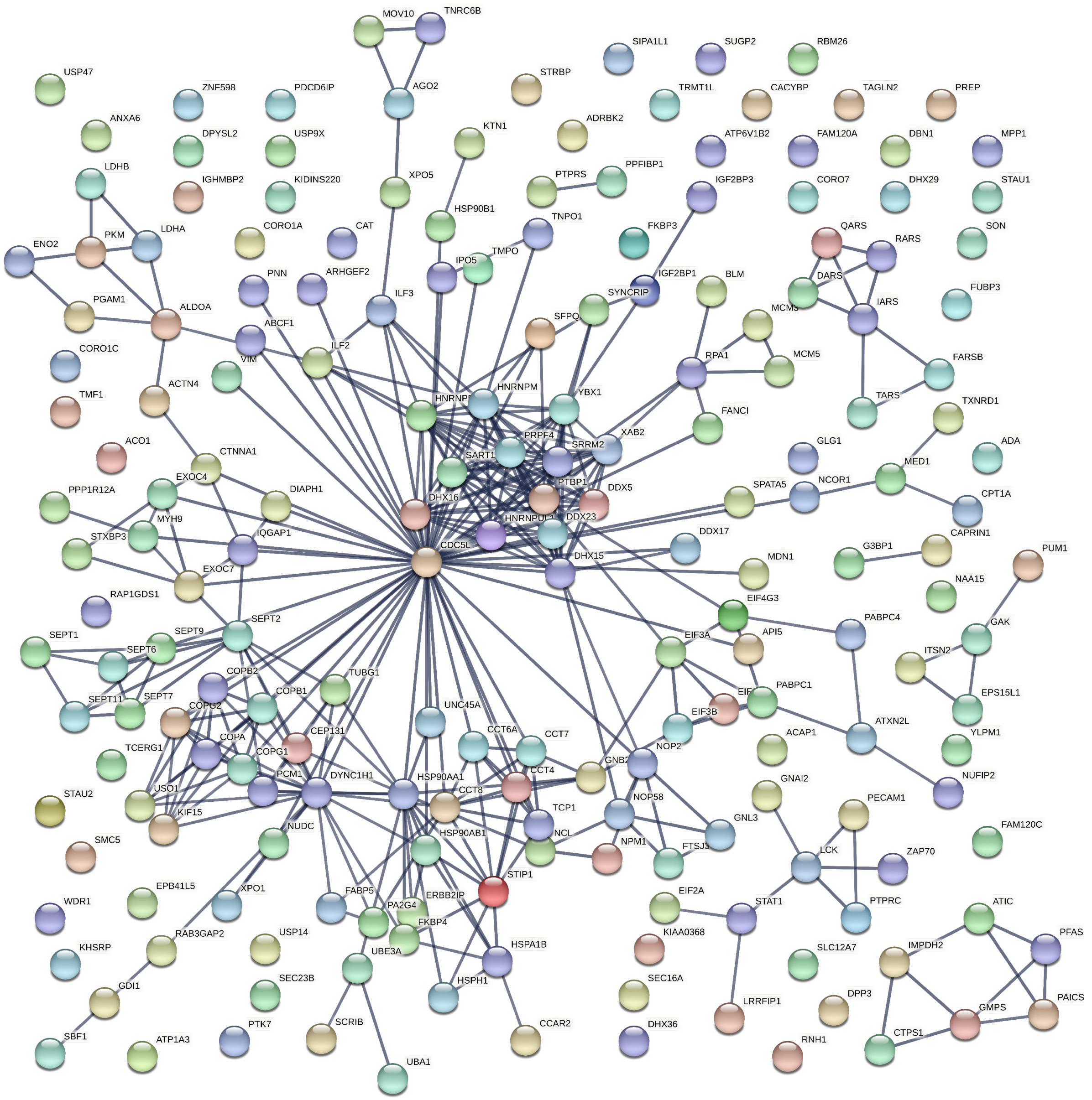

MOV10
TNRC6B
AGO2
SIPA1L1
SUGP2
RBM26
USP47
ZNF598
PDCD6IP
STRBP
TRMT1L
CACYBP
TAGLN2
PREP
ANXA6
DPYSL2
USP9X
KTN1
ADRBK2
ATP6V1B2
FAM120A
DBN1
MPP1
LDHB
IGHMBP2
KIDINS220
XPO5
PTPRS
PPFIBP1
IGF2BP3
CORO7
DHX29
STAU1
CORO1A
CAT
HSP90B1
TNPO1
FKBP3
QARS
SON
ENO2
PKM
LDHA
RARS
TMPO
IPO5
PNN
ARHGEF2
IGF2BP1
BLM
DARS
FUBP3
ILF3
SYNCRIP
IARS
PGAM1
ALDOA
ABCF1
SFPQ
CORO1C
ILF2
RPA1
MCM5
FARSB
VIM
TMF1
ACTN4
HNRNPM
YBX1
FANCI
TARS
TXNRD1
XAB2
ACO1
PRPF4
SRRM2
GLG1
ADA
CTNNA1
SART1
MED1
EXOC4
PTBP1
DDX5
SPATA5
NCOR1
CPT1A
PPP1R12A
DIAPH1
DHX16
DDX23
MYH9
DDX17
CAPRIN1
STXBP3
IQGAP1
CDC5L
DHX15
MDN1
G3BP1
PUM1
EXOC7
NAA15
RAP1GDS1
EIF4G3
SEPT2
PABPC4
GAK
SEPT1
SEPT9
EIF3A
API5
SEPT6
ITSN2
COPB2
TUBG1
PABPC1
SEPT11
SEPT7
COPB1
EIF3B
EPS15L1
COPG2
UNC45A
ATXN2L
CEP131
CCT6A
ACAP1
COPA
CCT7
YLPM1
TCERG1
NOP2
COPG1
CCT4
GNB2
PCM1
DYNC1H1
HSP90AA1
NUFIP2
USO1
CCT8
GNAI2
STAU2
KIF15
NOP58
PECAM1
TCP1
GNL3
NUDC
NCL
HSP90AB1
FAM120C
SMC5
NPM1
FTSJ3
LCK
ZAP70
STIP1
EPB41L5
XPO1
FABP5
ERBB2IP
EIF2A
WDR1
PA2G4
STAT1
PTPRC
ATIC
FKBP4
KIAA0368
PFAS
RAB3GAP2
USP14
UBE3A
HSPA1B
IMPDH2
KHSRP
SLC12A7
SEC23B
GDI1
HSPH1
SEC16A
LRRFIP1
DPP3
GMPS
PAICS
SBF1
PTK7
SCRIB
CCAR2
DHX36
RNH1
CTPS1
ATP1A3
UBA1

**Supplemental Figure S13:** STRING diagram of SVM-based BANDLE hits found only for WT replicates.

Network edges indicate confidence of protein connection at a minimum interaction score of 0.9.

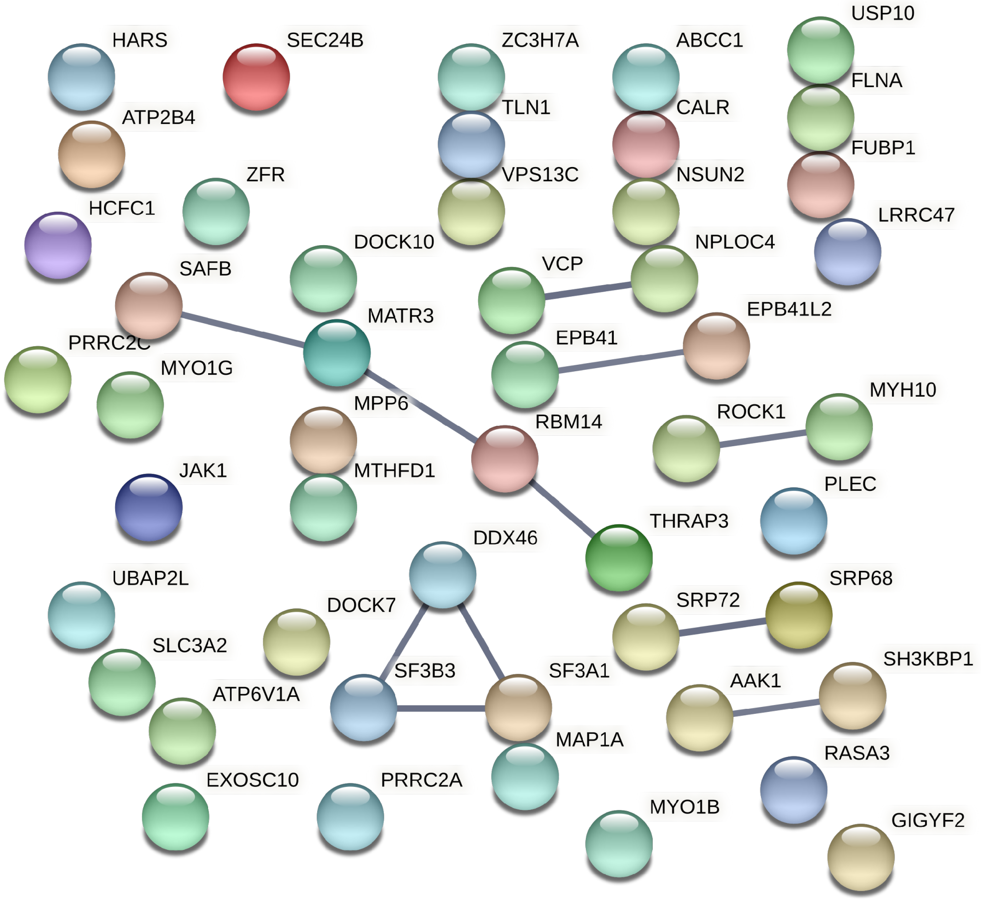
HARS
SEC24B
ZC3H7A
ABCC1
USP10
ATP2B4
TLN1
CALR
FLNA
FUBP1
ZFR
VPS13C
NSUN2
HCFC1
LRRC47
DOCK10
NPLOC4
SAFB
VCP
EPB41L2
MATR3
PRRC2C
EPB41
MYO1G
MYH10
MPP6
ROCK1
RBM14
JAK1
MTHFD1
PLEC
THRAP3
DDX46
UBAP2L
SRP68
DOCK7
SRP72
SLC3A2
SH3KBP1
SF3B3
SF3A1
AAK1
ATP6V1A
MAP1A
RASA3
EXOSC10
PRRC2A
MYO1B
GIGYF2

**Supplemental Figure S14:** STRING diagram of SVM-based BANDLE hits found for both WT and ΔNef replicates. Network edges indicate confidence of protein connection at a minimum interaction score of 0.9.

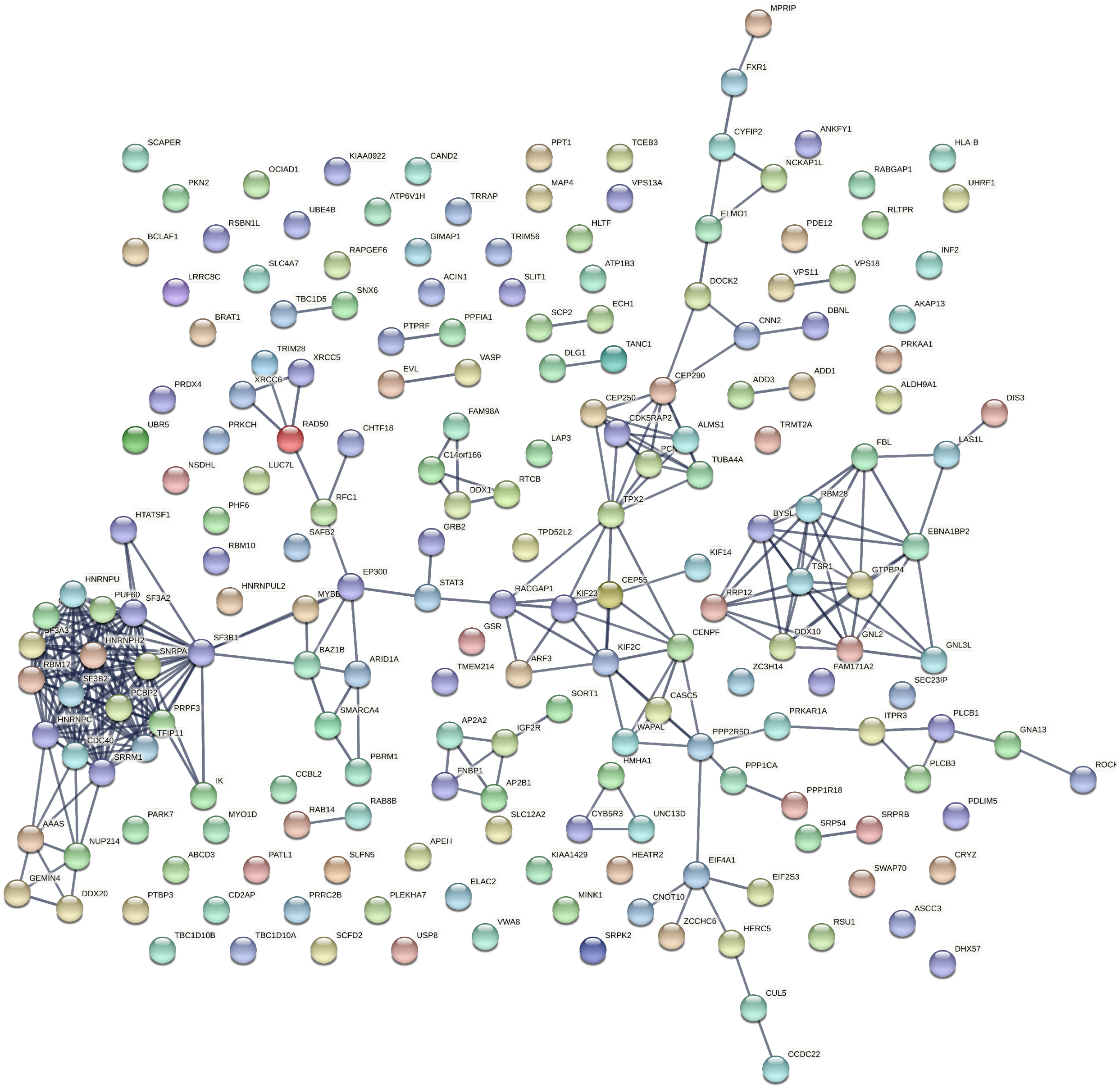
MPRIP
FXR1
CYFIP2
ANKFY1
NCKAP1L
HLA-B
SCAPER
KIAA0922
CAND2
PPT1
TCEB3
OCIAD1
PKN2
MAP4
VPS13A
RABGAP1
UHRF1
UBE4B
ATP6V1H
TRRAP
RSBN1L
HLTF
ELMO1
PDE12
RLTPR
BCLAF1
GIMAP1
TRIM56
RAPGEF6
ATP1B3
VPS11
VPS18
INF2
SLC4A7
ACIN1
SLIT1
LRRC8C
DOCK2
TBC1D5
SNX6
ECH1
AKAP13
SCP2
CNN2
DBNL
BRAT1
PTPRF
PPFIA1
TANC1
PRKAA1
TRIM28
XRCC5
DLG1
VASP
XRCC6
EVL
ADD1
CEP290
ADD3
PRDX4
CEP250
ALDH9A1
DIS3
RAD50
FAM98A
CDK5RAP2
UBR5
PRKCH
CHTF18
ALMS1
TRMT2A
LAS1L
LAP3
FBL
C14orf166
TUBA4A
NSDHL
LUC7L
RTCB
DDX1
RBM28
RFC1
TPX2
PHF6
BYSL
HTATSF1
SAFB2
TPD52L2
EBNA1BP2
GRB2
RBM10
KIF14
TSR1
EP300
CEP55
GTPBP4
HNRNPU
HNRNPUL2
STAT3
RACGAP1
KIF23
RRP12
PUF60
MYBB
SF3A2
SF3B1
SF3A3
HNRNPH2
SNRPA
GSR
CENPF
DDX10
GNL2
KIF2C
BAZ1B
ARID1A
GNL3L
ARF3
RBM17
TMEM214
ZC3H14
FAM171A2
SF3B2
SEC23IP
PCBP2
SORT1
CASC5
SMARCA4
PRPF3
PRKAR1A
HNRNPC
AP2A2
IGF2R
WAPAL
PPP2R5D
ITPR3
PLCB1
TFIP11
GNA13
CDC40
PBRM1
SRRM1
HMHA1
PPP1CA
PLCB3
ROCK
FNBP1
AP2B1
IK
CCBL2
PPP1R18
RAB14
RAB8B
CYB5R3
UNC13D
PDLIM5
PARK7
MYO1D
SLC12A2
SRP54
SRPRB
AAAS
NUP214
APEH
ABCD3
PATL1
SLFN5
KIAA1429
HEATR2
EIF4A1
SWAP70
CRYZ
GEMIN4
EIF2S3
ELAC2
DDX20
PTBP3
CD2AP
PRRC2B
PLEKHA7
MINK1
CNOT10
ASCC3
VWA8
ZCCHC6
HERC5
RSU1
TBC1D10B
TBC1D10A
SCFD2
USP8
SRPK2
DHX57
CUL5
CCDC22

**Supplemental Figure S15:** STRING diagram of SVM-based BANDLE hits found only for ΔNef replicates. Network edges indicate confidence of protein connection at a minimum interaction score of 0.9.
